# Protein-Folding Chaperones Predict Structure-Function Relationships and Cancer Risk in *BRCA1* Mutation Carriers

**DOI:** 10.1101/2023.09.14.557795

**Authors:** Brant Gracia, Patricia Montes, Angelica Maria Gutierrez, Banu Arun, Georgios Ioannis Karras

## Abstract

Identifying pathogenic mutations and predicting their impact on protein structure, function and phenotype remain major challenges in genome sciences. Protein-folding chaperones participate in structure-function relationships by facilitating the folding of protein variants encoded by mutant genes. Here, we utilize a high-throughput protein-protein interaction assay to test HSP70 and HSP90 chaperone interactions as predictors of pathogenicity for variants in the tumor suppressor BRCA1. Chaperones bind 77% of pathogenic BRCA1-BRCT variants, most of which engaged HSP70 more than HSP90. Remarkably, the magnitude of chaperone binding to variants is proportional to the degree of structural and phenotypic defect induced by *BRCA1* mutation. Quantitative chaperone interactions identified BRCA1-BRCT separation-of-function variants and hypomorphic alleles missed by pathogenicity prediction algorithms. Furthermore, increased chaperone binding signified greater cancer risk in human *BRCA1* carriers. Altogether, our study showcases the utility of chaperones as quantitative cellular biosensors of variant folding and phenotypic severity.

**HIGHLIGHTS:**

- Chaperones detect an abundance of pathogenic folding variants of BRCA1-BRCT.
- Degree of chaperone binding reflects severity of structural and phenotypic defect.
- Chaperones identify separation-of-function and hypomorphic variants.
- Chaperone interactions indicate penetrance and expressivity of *BRCA1* alleles.

## INTRODUCTION

Understanding how changes in genotype transform into phenotypes is paramount to predicting cancer risk and developing efficacious treatments in the clinic. In the era of personalized genomics, the number of variants of uncertain clinical significance (VUS) has grown faster than the rate of variant assignment^1–3^. In addition, there exists substantial discordance among leading commercial laboratories that classify variants and provide genetic testing services^4^. Meanwhile, relationships among variations and human health are archived into specialized databases, such as ClinVar, clinicians utilize to make informed decisions for patients carrying mutations^5^. However, the fraction of genetic variants classified as a practice guideline currently is miniscule (0.03%). Furthermore, estimating features of human heredity including reduced penetrance (likelihood of disease among individuals with similar genotypes), variable expressivity (differences in disease severity among patients with similar genotypes) and hypomorphism (partial loss of gene function due to mutation) remains exceptionally difficult to integrate into clinical practice^6–8^.

Protein folding is a fundamental process frequently disrupted by pathogenic mutations. Most proteins fold into specific three-dimensional structures encoded by the primary amino acid sequence^9–12^. Globular domains are the most well-characterized fundamental units of protein structure, which generally fold independently from the rest of the protein. However, folded domains tend to be marginally more stable than the unfolded state^13,14^. Thus, even small changes in DNA sequence can profoundly disrupt protein domain folding and function. Indeed, pathogenic missense mutations cluster within globular protein domains^12,15^. Although, not all pathogenic missense mutations affect protein structure (e.g., mutations in catalytic residues of enzymes or surface residues coordinating protein-partner interactions). However, the number of amino acids that contribute to domain folding greatly outnumbers interface or catalytic residues for most domains. Thus, it is likely that most pathogenic mutations within domains affect protein structure stability. Moreover, the marginal thermodynamic stability of protein structure can sensitize proteins to disease-associated environmental factors^16–19^. Hence, protein folding may be able to explain differences in the functional severity, penetrance and expressivity of human mutations.

Several prediction algorithms have been developed to infer the pathogenicity of mutations. These algorithms classify mutations by integrating available wild-type protein structures and sequence conservation from phylogenetic analyses^20–22^. Though facile, prediction algorithms vary substantially in their accuracy and face challenges in distinguishing benign from pathogenic mutations^23^. Recently, highly accurate structure prediction tools have been developed by combining fundamental principles of protein structure (e.g., secondary structure elements, hydrogen bonding networks and hydrophobic interactions) with co-variation analyses and machine learning^24^. However, these structure predictions breakdown when considering the impact of mutation on protein structure and function, likely because structure predictions are based on well-behaved “wild-type” proteins^25^. Furthermore, pathogenicity prediction algorithms do not account for influences originating from the environment, such as the crowdedness of the intracellular milieu where most disease-associated proteins are naturally found. Thus, a generalizable approach to measure changes in the folding of protein variants in a cellular context would greatly aid efforts to classify the clinical significance of genetic variation.

Protein-folding chaperones are highly conserved and abundant proteins that evolved to assist in the folding, maturation, localization and degradation of proteins within the cell^26–28^. Two of the most abundant cytosolic chaperones, HSP70 and HSP90, fold a large fraction of the human proteome including oncogenic kinases and tumor suppressor DNA repair proteins^29,30^. For this reason, quantitative HSP70 and HSP90 interactions with disease-relevant proteins have been useful for evaluating the impact of genetic and environmental factors on protein folding^31–34^. Previously, we utilized chaperone interactions to identify a new class of disease-associated mutations in *FANCA*^18^. The encoded FANCA protein variants interacted with chaperones in specific ways that correlated with the functional outcome of mutation, reflecting the unique substrate specificities of HSP70 and HSP90 chaperones. HSP70 binds stretches of amino acids that are highly hydrophobic and generally found in the core of globular domains^30^. In contrast, HSP90 preferentially binds partially folded, native-like proteins later during folding^35–37^. Because hydrophobic polypeptides are exposed at early stages of folding, we hypothesized that HSP70 preferentially binds protein variants adopting unfolded or misfolded conformations that are functionally compromised. Consistent with this hypothesis, cells carrying FANCA proteins variants that preferentially interact with HSP70 develop severe phenotypes equivalent to cells carrying *FANCA* gene deletions^18^. The generality of this hypothesis and whether chaperone-binding patterns can predict penetrance and expressivity of cancer mutations found in humans remains to be determined.

Tumor suppressor *BRCA1* is essential to life^38^. *BRCA1* inactivation by mutation increases the lifetime risk of developing breast and ovarian cancers by up to 7-8-fold in women^39^ and strongly predisposes individuals to pancreatic and prostate cancers^6,40^. The tumor suppressive role of *BRCA1* is attributable to the ability of the encoded BRCA1 protein to prevent a central hallmark of cancer, genome instability. In suppressing tumor development, BRCA1 serves as an enzyme and scaffold during DNA damage checkpoint regulation, activation of the transcriptional response to DNA damage and the repair of DNA double-strand breaks in homology-directed repair^41–44^. Genetic variations that disrupt BRCA1 protein structure or function drive genome instability phenotypes^45–47^. Thus, predicting the outcome of BRCA1 protein variants is critical to cancer risk management, diagnosis and treatment. Many independent approaches have evaluated *in vitro* the effects of *BRCA1* variations on protein function^49–53^, but whether data these approaches generate can integrate clinical features of human heredity or predict mutation-associated cancer risk remains unclear.

Here, we use a high-throughput and quantitative protein-chaperone interaction platform (LUMIER) to test if chaperone binding patterns predict structure-function relationships of BRCA1 variants. We find that most pathogenic variants in the BRCT domain of BRCA1 bind to chaperones in living cells, selectively to HSP70 over HSP90. We show that chaperones bind a variety of BRCA1-BRCT protein folding mutants, and the pattern of chaperone binding reflects specific structural and functional disruptions induced by each variant. Remarkably, a class of variants that bind chaperones retained partial structure and function. Moreover, we show that the magnitude of chaperone binding is proportional to the functional severity of the corresponding allele, with hypomorphic alleles exhibiting intermediate chaperone binding to the encoded protein variant. Furthermore, patients carrying germline BRCA1 variants that bind chaperones exhibit increased disease penetrance and earlier age of cancer onset. Altogether, these results suggest that chaperone interactions to protein variants carried in people can serve as a proxy for disease severity. We propose parsing variations in disease-relevant genes based on the chaperone binding pattern of the encoded variant protein as a powerful strategy with broad applicability for risk stratification and precision medicine applications.

## RESULTS

### Most Pathogenic BRCA1-BRCT Variants Bind Chaperones

A large fraction of BRCA1 missense variants in ClinVar are of uncertain/conflicting clinical significance (Figure 1A, n=3,591). 93% of pathogenic missense mutations are within the N-terminal RING domain or the C-terminal BRCT domain (Figure 1B). The BRCA1-RING domain requires heterodimerization with the BARD1-RING domain to reconstitute a functional E3 ubiquitin ligase^53^. In contrast, the BRCA1-BRCT includes two tandem BRCT subdomains that interact to form a hydrophobic groove that recognizes specific phosphorylated peptides (pSXXF) in DNA repair proteins critical for tumor suppression by BRCA1 (Figure 1C)^43,54^. We expressed the BRCA1-RING and BRCA1-BRCT domains fused to a 3xFLAG tag and measured chaperone interactions in living cells using a high-throughput and quantitative coimmunoprecipitation assay (Figure 1D, STAR Methods LUMIER with BACON)^31,55^. For this, we used NanoLuc luciferase-tagged chaperones because they produced higher signal to noise as compared to the published *Renilla* luciferase system (Figure S1A). In contrast to established HSP70 and HSP90 clients and co-chaperones, the wild-type BRCA1-BRCT domain did not bind to either chaperone significantly in our assay (Figures S1B-C, HSP70 mean±st.dev. z-score=1.4±0.8, HSP90 mean±st.dev. z-score=-0.2±1.4). On the other hand, the wild-type BRCA1-RING domain strongly interacted with HSP70 and did not bind HSP90 (Figure S1D, HSP70 z-score=5.0±0.9, HSP90 z-score=0.7±0.8). These results suggest that the wild-type BRCA1-BRCT is well-folded in cells because the domain quickly folds and releases from chaperones, unlike the BRCA1-RING domain folding which relies on BARD1-RING binding^42,53,56^.

**Figure 1.**
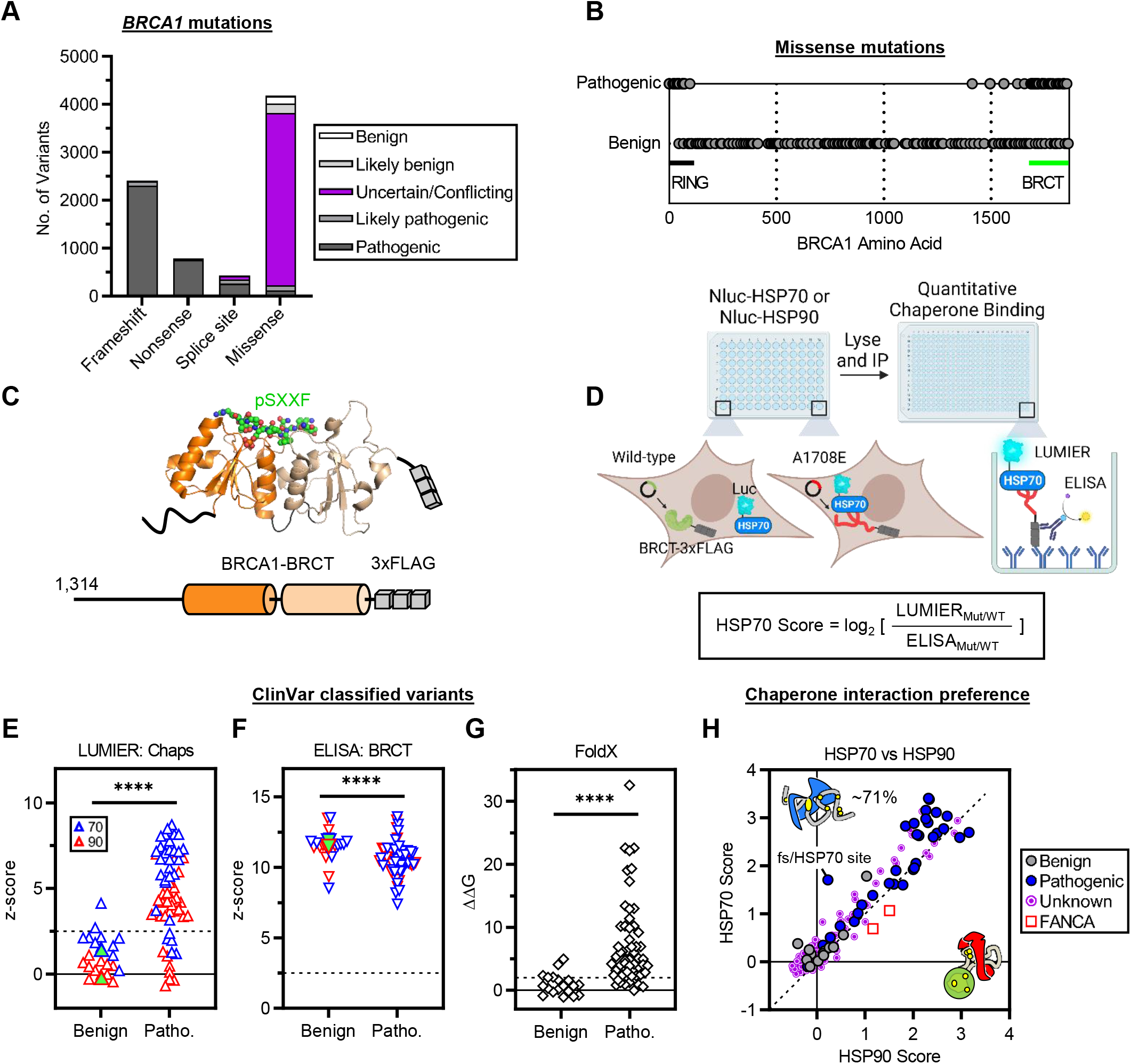
Most pathogenic BRCA1-BRCT variants bind chaperones. (A, B) *BRCA1* mutations catalogued in ClinVar^5^. (A) Clinical significance of *BRCA1* mutations grouped by mutation type. (B) Missense mutations catalogued in ClinVar with at least a ‘two’ gold star review status (n=285). ClinVar accessed 4/27/23. (C) C-terminal BRCA1 3x-FLAG tagged truncation construct. The two BRCT subdomains are colored (orange and wheat). pSXXF denotes BRCT binding phosphopeptide (BACH1), PDB: 1t29. (D) LUMIER with BACON approach to quantify chaperone interactions in HEK293T cells. Variant HSP interaction scores calculated by log_2_ transforming the data followed by subtracting LUMIER by ELISA values for variant (mut) and wild-type (WT), separately, and then subtracting the variant value by the wild-type value. For simplicity, the equation shown compresses the calculation into a single log_2_. Diagram created with BioRender.com. (E) Chaperone binding to benign and pathogenic BRCA1-BRCT variants from ClinVar (including ‘likely’ classifications). Z-scores shown relative to negative control wells. Dashed lines indicate threshold for statistically significant signal (z-score>2.5). Wild-type BRCA1-BRCT values shown as green filled symbols. (F) BRCA1-BRCT variant abundance detected by ELISA. (G) Variant ΔΔG values (kcal/mol) predicted by FoldX relative to the wild-type (ΔΔG=0). Dashed line shows threshold for structure disruption (ΔΔG>2). (H) HSP70/HSP90 interaction preference. BRCA1-BRCT and FANCA variants are normalized to their respective wild-type protein. Diagonal shows identity line. Unknown group includes variants annotated as uncertain significance, no significance provided, conflicting interpretations, and ‘one’ gold star review status. FANCA variants R880Q and I939S and frameshifted (fs) variant with additional HSP70 site shown for comparison. Statistical significance was determined using two-tailed Mann-Whitney t-test (E, F and G). ****p≤0.0001. Data are represented as mean from at least two independent experiments.

We cloned a library of clinical variants in the BRCA1-BRCT domain (ClinVar: benign, n=11; pathogenic, n=31; unknown, n=118), BRCA1-RING domain (benign, n=2; pathogenic, n=14; unknown n=10) and control variants not part of either domain (N-terminal region, n=9; C-terminal region, n=10), and measured the binding of each variant to HSP70 and HSP90. Control variants outside of the RING or BRCT domain did not change binding, except for a frameshifting variant that introduced a new HSP70 binding site not present in the wild-type open reading frame. Interestingly, BRCA1-BRCT variants exhibited dramatic increases in chaperone binding that varied drastically depending on the variant. The increases in chaperone binding to different BRCA1-BRCT variants were highly reproducible between independent experiments (Figure S1E, HSP70 R^2^=0.91 and HSP90 R^2^=0.92, *p*≤0.0001). On the other hand, pathogenic and benign/unknown BRCA1-RING domain variants did not significantly alter HSP70, in line with the wild-type BRCA1-RING domain not folding properly in our assay (Figure S1D). Thus, we focused on chaperone interaction patterns to disease-associated BRCA1-BRCT variants.

Importantly, ∼77% (24 out of 31) of pathogenic BRCA1-BRCT variants in our ClinVar library significantly interacted with both chaperones (Figure 1E, z-score>2.5). In contrast, ∼82% (9 out of 11) of benign variants did not interact with either chaperone as compared to the wild-type BRCA1-BRCT (*p*≤0.0001). Because chaperones are involved in the degradation of misfolded proteins^57,58^, we tested if increased chaperone binding correlates with changes in protein abundance. We used an ELISA approach to quantify BRCA1-BRCT variant abundance after pull-down and obtained ELISA signals that were significantly above background and highly reproducible (Figure S1F, R^2^=0.94, *p*≤0.0001). Using ELISA, we observed that pathogenic BRCA1-BRCT variants were ∼2-3-fold less abundant as compared to benign variants and the wild-type (Figure 1F, *p*≤0.0001), suggesting that pathogenic variants are less stable and degraded in cells. Furthermore, increased chaperone binding and decreased protein abundance for pathogenic BRCA1-BRCT variants were reproduced using conventional coimmunoprecipitation and Western Blotting against endogenous chaperones (Figure S1G). LUMIER interactions were also validated in 3 different BRCT-containing BRCA1 fragments fused to different tags (Figure S1H). Lastly, *in silico* predictions using FoldX^20^ suggest that 86% of pathogenic BRCT missense variants from ClinVar increase the Gibbs free energy of folding as compared to benign variants (Figure 1G, ΔΔG>2 kcal/mol, *p*≤0.0001). Taken together, these results suggest that most pathogenic BRCA1-BRCT missense variants bind chaperones and are unstable in cells.

Next, we generated variant HSP interaction scores relative to wild-type by normalizing for differences in ELISA (Figure 1D) and evaluated the HSP70/HSP90 chaperone binding pattern of pathogenic BRCA1-BRCT variants. We observed that pathogenic BRCA1-BRCT variants are predominantly engaged by HSP70 over HSP90 (∼71%), especially when compared with HSP90-engaged FANCA variants that retain protein function^18^ (Figure 1H). In addition, ∼29% of unknown significance (conflicting, uncertain, not provided, one star review status) BRCA1-BRCT variants bind to chaperones (Figure 1H, HSP70/HSP90 scores>0.5) suggesting chaperone interaction patterns can help assign clinical significance to human BRCA1-BRCT variants. Notably, chaperone binding varied drastically (1.5 to 11-fold) between BRCA1-BRCT variants, substantially more than the increases in chaperone binding previously observed for FANCA variants^18^. The broad diversity of chaperone binding patterns observed suggest that chaperones detect different types of BRCA1-BRCT folding variants.

### Chaperone Biosensors Detect a Diversity of Protein Folding Variants

A possible explanation for the increased binding of BRCA1-BRCT variants to HSP70 is that the variants we tested increase the number of HSP70 binding motifs, thus changing the binding stoichiometry of HSP70. Contrary to this scenario, we observed no statistically significant difference between variants predicted to harbor more/larger DnaK binding sites (bacterial HSP70) than the wild-type BRCA1-BRCT as compared to variants with unchanged or fewer/smaller sites (Figures S2A-B). Instead, chaperone binding to BRCA1-BRCT variants appears to reflect changes in domain thermodynamic stability. To test this, we compared BRCA1-BRCT variant chaperone binding with a previously published protease sensitivity assay^46,47,59^. In this assay, variants that have structural disruptions relative to wild-type exhibit increased sensitivity to protease cleavage. We observed increased HSP70 binding and decreased bait abundance for BRCA1-BRCT variants that are highly sensitive to protease digestion relative to control variants (Figure S2C, *p*≤0.0001). The data suggest that increased chaperone binding to BRCA1-BRCT variants generally reflects changes in thermodynamic stability. To closely examine the relationship between chaperone binding and thermodynamic stability, we rationally designed variants (n=49) that disrupt fundamental elements of protein structure and measured chaperone binding to each engineered variant. For this, we focused on secondary structure elements, side chain hydrogen bonds and hydrophobic interactions.

We first attempted to disrupt secondary structure elements observed in the BRCA1-BRCT crystal structure. Proline variants are typically not tolerated in α-helix secondary structures^60^. FoldX predicts strong structural disruptions when helical residues are mutated to proline as compared to alanine (Figure S2D, *p*≤0.0001). Additionally, proline is the most frequent pathogenic mutation observed for BRCA1-BRCT α-helical residues in ClinVar (5 out of 19). As expected, we observed increased chaperone binding to the BRCA1-BRCT when prolines were swapped in at α-helices for 4 out of 4 designed variants (Figure 2A: V1665P, Y1666P, T1720P, D1840P). Similarly, charged variants introduced at β-sheets are frequently pathogenic (9 out of 20) and FoldX predicts they disrupt the BRCA1-BRCT structure. We observed increased chaperone binding to the BRCA1-BRCT domain when glutamates were swapped in at each BRCT subdomain β-sheet for 2 out of 2 designed variants (Figure 2A, V1687E and I1807E). Interestingly, variants designed to disrupt α-helices or β-sheets were engaged predominantly by HSP70 over HSP90 (e.g., I1807E binds 2.1-fold more to HSP70 *vs* HSP90).

**Figure 2.**
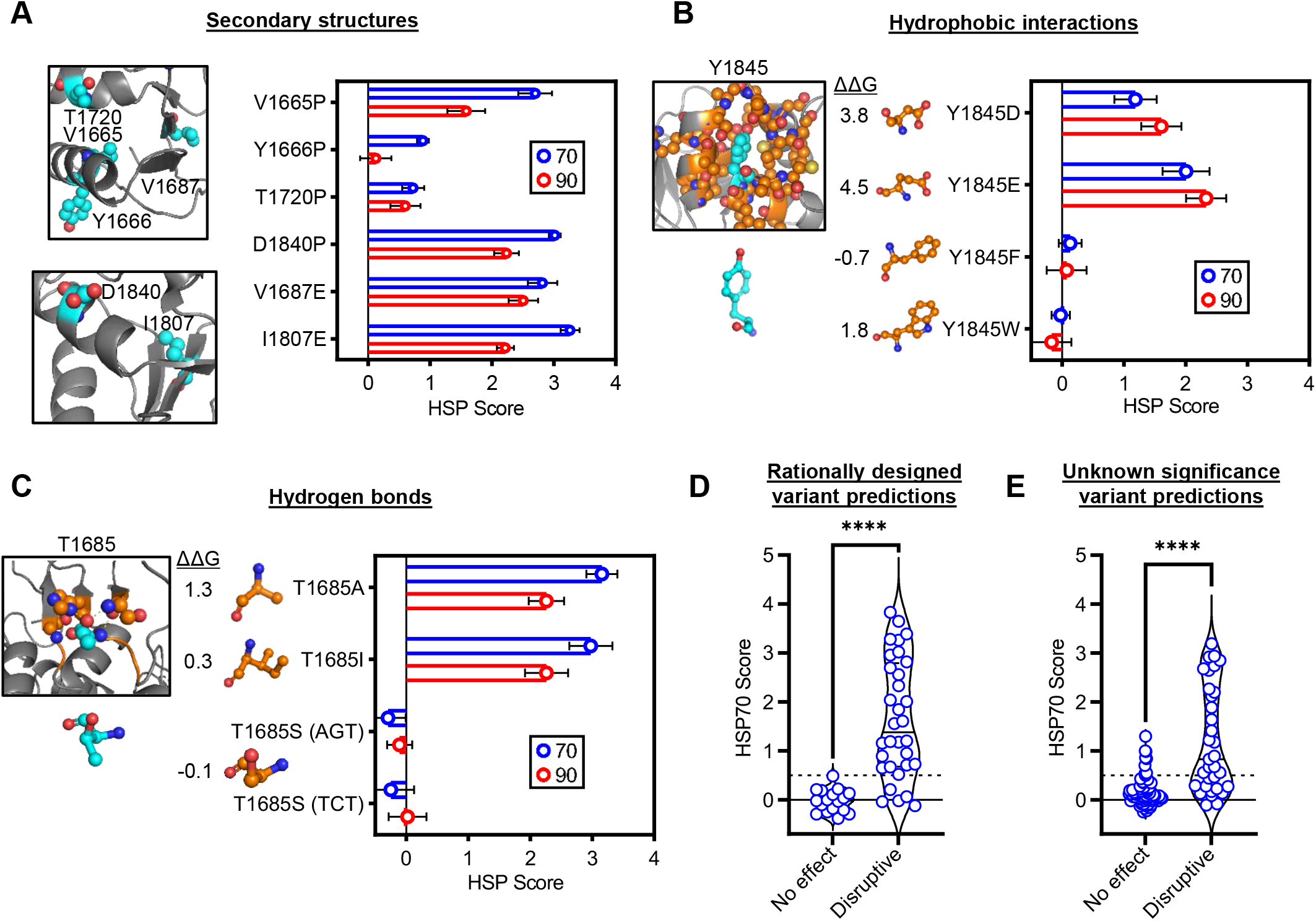
Chaperone biosensors detect a diversity of protein folding variants. (A) Variants targeting secondary structure elements. (B) Y1845 variants to disrupt or support hydrophobic interactions. FoldX ΔΔG predictions for each variant are shown. (C) T1685 variants to disrupt or support side chain hydrogen bonding. Two different T1685S codons were tested because previous functional data showed differential effects for these codons^50^. (D, E) Chaperone binding predictions to BRCA1-BRCT variant library. Dashed line shows cutoff for binding (HSP70 score>0.5). Unknown variant predictions from ClinVar excludes variants previously characterized by protease sensitivity^47^. Statistical significance was determined using two-tailed Mann-Whitney t-test. ****p≤0.0001. Data are represented as mean±standard deviation from at least two independent experiments.

Hydrophobic interactions in the core of folded domains are critical for protein folding and stability^61^. We rationally designed 11 variants to target hydrophobic interactions observed in the BRCA1-BRCT structure. We observed increased chaperone binding when charged amino acids were swapped in at buried residues that form hydrophobic interactions (Figures 2B and S2E: Y1845D, Y1845E, M1783R, M1783K), suggesting charged variants at these positions disrupt hydrophobic interactions. In contrast, biochemically compatible variants did not increase chaperone binding (e.g., Y1845F, Y1845W) or weakly increased chaperone binding (e.g., M1783A). Overall, we observed greater chaperone binding to variants in our library that mutate buried residues relative to surface residues (Figures S2F, buried: mean HSP70=1.5, intermediate/surface: mean HSP70=0.4, *p*≤0.0001). Altogether, our findings demonstrate that disrupting critical structural elements of BRCA1-BRCT domain architecture drastically increase chaperone binding, and greater HSP70 binding reflects strongly destabilized BRCA1-BRCT variants.

Next, we tested if chaperone interactions can detect disruptions to hydrogen bonding networks involving residue side chains. For this, we targeted residues on the surface of the BRCA1-BRCT so that loss of hydrogen bonds would not compromise other structural elements. T1685 forms side chain hydrogen bonds with the backbone amide of S1651 and the side chain of H1686 on the surface of the BRCA1-BRCT (Figure 2C). Two pathogenic ClinVar variants from our library block these hydrogen bonds (T1685A and T1685I), and both variants were predominantly engaged by HSP70 (T1685A: HSP70=3.2±0.3). T1685A preferential binding to HSP70 did not reflect increased number of HSP70 binding sites because T1685 variants do not change the number of HSP70 binding sites predicted computationally. Thus, we tested if hydrogen bonds formed by the side chain of T1685 are important for folding by engineering a variant that preserves hydrogen bonding. Indeed, mutating T1685 to serine (T1685S) did not increase chaperone binding in two different codon backgrounds tested (T1685S (TCT): HSP70=-0.2±0.4). Hence, the hydroxyl side-chain of T1685 coordinates a network of hydrogen bonds at the BRCA1-BRCT surface that stabilize the structure of the domain. We tested if additional surface residues function similarly by designing 10 surface variants to target side chain hydrogen bonds. We identified 9 wild-type residues that increase chaperone binding when the variant disrupts side chain hydrogen bonds (Figure S2G), suggesting the wild-type side chain hydrogen bonds are surface determinants of BRCA1-BRCT folding. Notably, for these surface variants that disrupt side chain hydrogen bonding, FoldX predicted weak effects on protein stability (mean ΔΔG=1.2±0.9). We conclude that chaperones can detect BRCA1-BRCT variants that disrupt side chain hydrogen bonding networks, even when the variants impact on the folded structure is subtle.

There are examples where the effect of mutation on chaperone binding is complex and difficult to explain quantitatively. Mutating charged residues at the N or C termini of certain α-helices can disrupt protein structure by influencing helix dipole moments^62,63^. We mutated charged residues at the termini of α-helix 1 and observed chaperone binding specifically to E1661K (Figure S2H, HSP70=1.6±0.2) as compared to other charge swapping variants which did not increase chaperone binding (E1660K, R1670E and K1671E). However, residue E1661 also forms hydrogen bonds via its side chain with K1690, T1658 and the pSXXF mimic which together may stabilize the BRCA1-BRCT structure. Indeed, disruption of this hydrogen bonding network by mutation of E1661 to alanine (E1661A) moderately increased HSP70 binding, while a variant compatible with hydrogen bonding, E1661Q, showed wild-type-like chaperone binding. Notably, E1661A binds chaperones ∼2-fold less strongly than E1661K, suggesting the E1661K variant disrupts folding by other ways than mere disruption of hydrogen bonding. We also targeted residue T1658 which hydrogen bonds with the peptide backbone of E1660 and E1661 and caps the N-terminus of α-helix 1. However, disrupting these hydrogen bonds did not affect chaperone binding (Figure S2H, T1658A, T1658N and T1658S), suggesting they are not important for domain folding, represent BRCA1-BRCT crystallization artifacts, or loss of stabilization at T1658 is rescued by neighboring residues (e.g. E1660 could theoretically stabilize the helix in the absence of T1658). Although the mechanism of BRCA1-BRCT structure disruption by E1661K is unclear, the increased chaperone binding of the variant suggests it is clinically relevant. Indeed, this variant has been observed in a human cancer patient (skin cancer, non-melanoma) and is annotated as conflicting interpretations of pathogenicity in ClinVar.

Another set of variants difficult to interpret involved residues within the hydrophobic interface between the BRCA1-BRCT subdomains. Two pathogenic variants in our library (A1708E and G1706E) introduce a charge within this groove and bind strongly to chaperones (Figure S2I). We generated 8 additional variants in this region to preserve or disrupt the local biochemical environment. Interestingly, these mutations increased chaperone binding to a more variable extent. Variants that preserve hydrophobicity or suitably accommodate the steric constraints of the pocket interacted with chaperones only moderately (L1705I, L1705V and G1706A) as compared to variants that introduce a charge (L1705K, L1705R, L1705D, A1708R and F1704D). Notably, three of these variants that introduce a charged amino acid into the interface between BRCT subdomains were predominantly engaged by HSP90 (F1704D, G1706E and A1708E), suggesting the variant disrupts subdomain interactions more subtly than variants that selectively engage HSP70. Hence, chaperones bind variants in different ways that reflect the structural effect of the specific mutation.

Altogether, the results in this section show that chaperones bind variants that disrupt protein structure. For our rationally designed library, we predicted 17 out of 17 (100%) variants would have no effect on structure (no increase in HSP70 binding) using simple energetic principles of thermodynamics (Figure 2D, HSP70 score cutoff=0.5). Analogously, 27 out of 32 (∼84%) variants expected to disrupt structure interacted with HSP70, amounting to a total accuracy of ∼90% for rationally designed variants. For unknown clinical significance variants from ClinVar, 67 out of 76 (∼88%) variants predicted to not affect structure did not bind HSP70 (Figure 2E). Similarly, 21 out of 33 (∼64%) variants expected to disrupt structure interacted with HSP70, amounting to a total accuracy of ∼81% for unknown ClinVar variants in our library. In conclusion, increased chaperone binding can help identify BRCA1-BRCT variants that disrupt structure even when the mechanism of disruption is not known.

### The Degree of Chaperone Binding is Proportional to the Severity of *BRCA1* Mutation

Because BRCT structure defines its function, we asked whether the degree of increased chaperone binding to variants reflects fractional loss of BRCA1-BRCT function. To determine this fraction, we first quantified the dynamic range of BRCA1-BRCT structure disruption as detected by chaperone binding. We quantified maximal chaperone binding by generating 13 double BRCA1-BRCT variants, combining single variants that each bind chaperones to differing degrees (e.g., weak paired with intermediate, strong paired with strong). Combining single variants that do not bind chaperones with other variants had no effect on HSP70 binding (Figure S3A, e.g. F1675L/D1778G, F1695L/M1775K and F1695L/V1809F). In contrast, combining two variants that each interacted moderately with HSP70 increased chaperone binding to the double variant (S1655F/M1775K, ∼4.3-fold increased binding relative to single variants). Strikingly, combining two strongly HSP70 interacting variants only weakly increased chaperone binding to the double variant (e.g., C1697R/D1840P and V1687E/G1788V, ∼1.5-fold increase binding relative to single variants). The increase in chaperone binding observed for these double variants was substantially smaller than predicted by additivity of the single variants (Figure S3A, ∼4.1-fold less than expected). Indeed, we found that HSP70 binding did not exceed an upper limit of ∼14-fold greater chaperone binding to variants as compared to the wild-type BRCA1-BRCT. FoldX was unable to predict this upper limit, as stability values observed for the double variants were the same as predicted by additivity. These results suggest that maximal chaperone binding in our assay reflects complete structural disruption of the BRCA1-BRCT variant.

Below the upper limit of chaperone binding, we observed a strong positive correlation between the degree of chaperone binding to each BRCA1-BRCT missense variant and the predicted increase in Gibbs free energy, as determined by FoldX (Figure 3A, FoldX *vs* HSP70: R^2^=0.63, *p*≤0.0001). This correlation was reproduced when using empirically derived stability measures for 18 recombinant BRCA1-BRCT variants purified from *E. coli* (Figure 3B, R^2^=0.78, *p*≤0.0001)^64^. Thus, increased chaperone binding to BRCA1-BRCT variants reflects loss of BRCT folding. In addition, 10 variants that were insoluble and could not be purified in *E. coli* interacted strongly with HSP70 (mean HSP70 score=2.7±0.7). Similar results were obtained for HSP90 binding scores (Figures S3B-C). These findings suggest that strong HSP70 binding to variants reflects severely disrupted BRCA1-BRCT domain structure as compared to intermediate chaperone binding to BRCA1-BRCT variants which generally reflects partial defects in protein folding.

**Figure 3.**
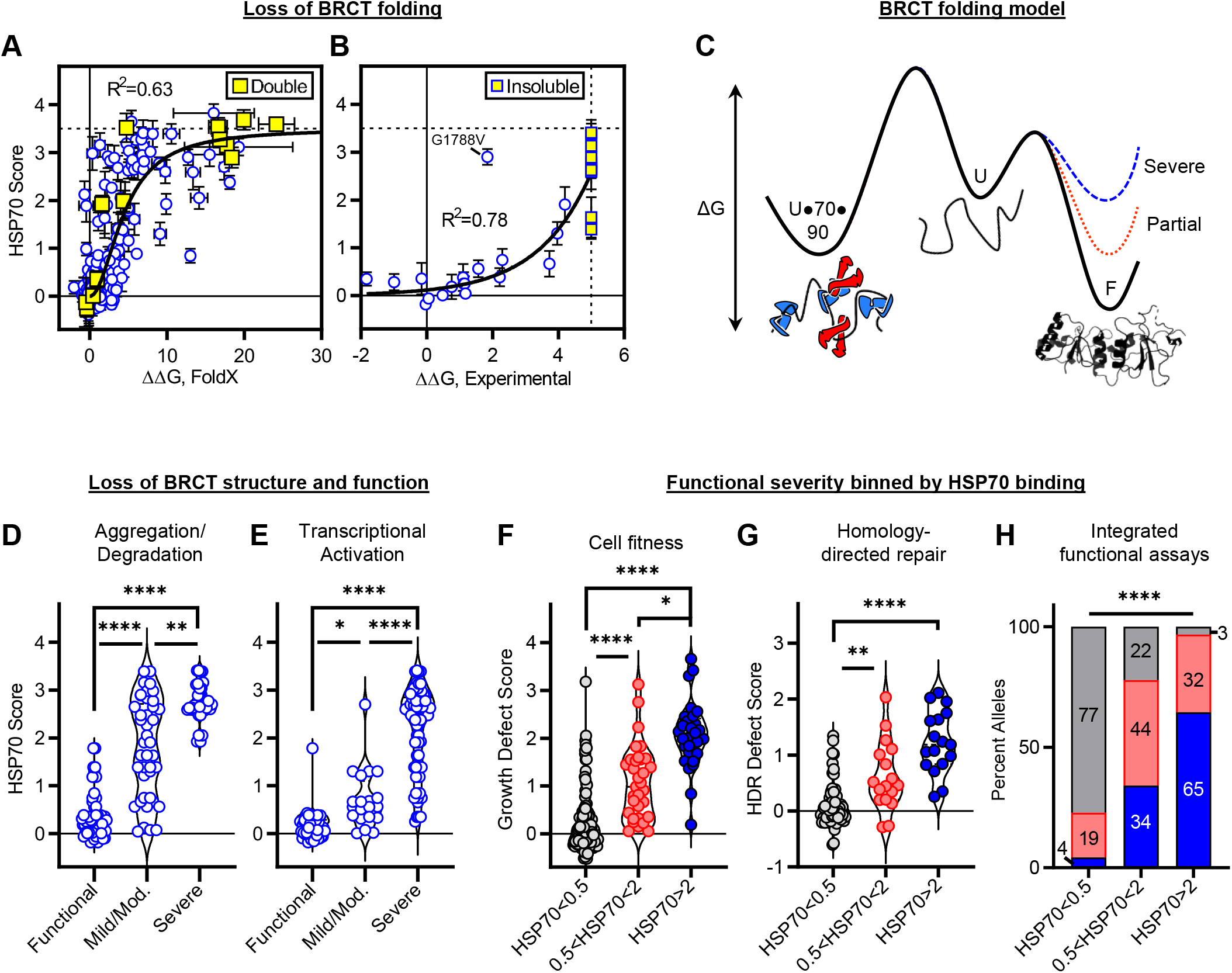
The degree of chaperone binding is proportional to the severity of BRCA1 mutation. (A, B) HSP70 binding to BRCA1-BRCT variants correlated with protein folding stability. Horizontal dashed line is the average of four double variants that combine strongly chaperone bound single variants. Vertical dashed line in (B) reflects an upper limit on stability measurements due to variant insolubility. FoldX data in (A) fit to the hill equation with hill coefficient set to 2 to account for HSP70 dimerization. Experimental data in (B) fit to a single exponential growth (excluding insoluble variants). G1788V not fit because stability measurements on this variant were previously reported as contradictory^64^. (C) Folding free energy profiles illustrating increased chaperone binding to BRCT variants that severely disrupt (blue) or partially disrupt (red) the folded state. Variant effects shown exclusively to the folded ground state (‘F’) relative to the unfolded state (‘U’) and unfolded chaperone bound state (‘U●70●90’). (D, E) HSP70 binding to BRCA1-BRCT variants binned by the functional effect measured in aggregation/degradation and transcriptional activation assays^65^. The same variant may appear more than once if the functional assay or reporting publication were different. (F, G) BRCA1 variant function in HAP1^50^ (cell fitness) or HeLa^52^ (homology-directed repair) cells binned by the degree of HSP70 binding. (H) Integrated functional data binned by the magnitude of HSP70 binding. Each variant was assigned a functional severity class using the mode phenotype observed from all available assays (functional, mild, moderate, severe, or ambiguous if the severity was multi-modal)^65^. Gray, functional; Red, mild/mod./ambiguous; Blue, severe. Residues that confer separation-of-function phenotypes in HAP1 and HeLa cells (n=9 variants) were filtered out. Statistical significance was determined using Kruskal-Wallis ANOVA test (D, E, F and G) or Chi-squared test (H). ****p≤0.0001, **p≤0.01, *p≤0.05. Data are represented as mean±standard deviation from at least two independent experiments.

We propose a model whereby the wild-type BRCA1-BRCT folds rapidly and favorably to the native structure, preventing subsequent rounds of chaperone binding to unfolded conformations (Figure 3C, black curve). Variants that severely disrupt the folded BRCA1-BRCT structure shift the folding equilibrium to the unfolded state, allowing for favorable accumulation of chaperone binding (blue curve). In contrast, partially disrupted BRCA1-BRCT variants spend less time in the unfolded conformation than severely disrupted variants and thus bind moderately to chaperones (red curve). In summary, the model reflects that increased chaperone binding to variants correlates with loss of BRCA1-BRCT folding. However, it remained unclear whether the degree of chaperone binding correlates with loss of BRCA1 protein function.

To test if the degree of chaperone binding is proportional to the functional (phenotypic) severity of the corresponding BRCA1 protein variant, we collected data from the NeXtProt server^65^, a rich resource that catalogues the functional severity of BRCA1 variants measured in many different assays and laboratories (n=573 unique entries for variants in our library). As expected, when the aggregation/degradation propensity of the variants increased so did chaperone binding (Figure 3D, mild/mod. *vs* functional or severe *p*≤0.009). Importantly, the same correlation was reproduced across diverse functional assays even in the context of full-length BRCA1, including transcriptional activation, cell localization and cell viability (Figures 3E and S3D-E). The results suggest that the degree of chaperone binding to BRCA1-BRCT variants reflects the functional severity of BRCA1 mutants in cells.

To further test the relationship between chaperone binding and functional severity of BRCA1 mutations in cells, we collected functional data from two independent high-throughput BRCA1 functional assays. We binned the data by the degree of HSP70 binding and investigated the growth defect score induced by BRCA1 missense mutation at the endogenous *BRCA1* locus in HAP1 cells^50^. Similar to results obtained using data from the NeXtProt server, we observed gradated functional defects that increased progressively with the magnitude of chaperone binding. BRCA1-BRCT variants weakly bound by HSP70 were predominantly functional in the context of the full-length BRCA1 protein in HAP1 cells, as compared to variants that strongly bind HSP70 which were severely defective for cell growth phenotypes (Figure 3F, HSP70<0.5: growth defect score mean=0.25±0.64, HSP70>2: growth defect score mean=2.1±0.7, *p*≤0.0001). Moreover, BRCA1-BRCT variants moderately bound by HSP70 conferred intermediate effects on phenotype, which were significantly different from weak and strong HSP70 binding variant groups (0.5<HSP70<2: growth defect score mean=1.1±0.8, *p*≤0.005). We observed a similar relationship between BRCA1-BRCT variant binding to chaperones and the effect of *BRCA1* mutation on homology-directed repair efficiency (HDR) in HeLa cells^52^ (Figure 3G). FoldX predictions recapitulated the above results (Figures S3F-G). These data further suggest that the degree of chaperone binding to BRCA1-BRCT variants is proportional to the degree of structural disruption and phenotypic severity of full-length mutant BRCA1 in cells. Indeed, we observed a strong positive linear correlation between functional defect induced by BRCA1 mutation and the magnitude of HSP70 binding to the variant (Figures S3H-I, HAP1: R^2^=0.52, *p*≤0.0001; HeLa: R^2^=0.49, *p*≤0.0001).

Although increased chaperone binding correctly identifies most structure disrupting BRCA1-BRCT variants, it fails to detect certain deleterious mutations. Indeed, ∼24% of variants in our library (16 out of 67) compromised BRCA1 function in HAP1 without increasing chaperone binding (∼35% in HeLa HDR assay), which suggests the variant defect does not involve structural disruption (Figures S3H and S3I). We delineate these variants as separation-of-function because the defect in BRCA1-BRCT function is separable from domain structure disruption. By comparing the position of residues in the BRCT crystal structure, we observed that ∼44% (7 out of 16) of separation-of-function variants (HAP1) are near the pSXXF (∼33% in HDR assay), suggesting these variants disrupt pSXXF binding and not folding (Figure S3J). Using this approach, we also observed 7 separation-of-function variants in HAP1 (2 variants in HeLa HDR assay) on the surface of the BRCT that are not near the pSXXF, suggesting these variants disrupt protein-partner interactions through a mechanism independent of the pSXXF. The remaining separation-of-function variants reside outside of the BRCT domain crystal structure suggesting disruption of other protein-partner motifs. For BRCT variants, we also observed that different variants of the same residue exhibited distinct effects, often involving complex consequences on domain folding and protein-partner interactions. Strikingly, well characterized contact variants^47,64,66^ interacted moderately with chaperones (e.g., S1655F, R1699W, M1775K, and M1775R), suggesting they partially disrupt domain folding. We conclude that integrating chaperone interactions with functional data can help distinguish true separation-of-function mutations from complex mutations that disrupt contacts with BRCA1 partners and destabilize the BRCT domain.

Altogether, our findings highlight the ability of chaperones to gauge the functional severity of *BRCA1* mutations measured in cell-based assays. After removing 9 separation-of-function mutations and aggregating all functional data from the NeXtProt server, we find that 77% of BRCA1-BRCT variants that bind weakly to HSP70 are functional, 44% of variants that bind moderately confer mild/moderate defects and 65% of variants that bind strongly exhibit severe functional defects (Figure 3H, *p*≤0.0001). We conclude that the degree of chaperone binding to BRCA1-BRCT variants is generally proportional to the phenotypic severity of the corresponding BRCA1 variant. Thus, quantitative chaperone interactions can be used to identify allele classes in cells including functional, severe, and hypomorphic (partial loss of function).

### Chaperones identify diverse classes of pathogenic *BRCA1* variations in people

We compared the frequency of variants that bind moderately to chaperones across diverse clinical and population studies to ascertain the abundance of hypomorphic BRCA1 variants in humans. BRCA1-BRCT variants that bind moderately to chaperones are abundant among annotated pathogenic variants in ClinVar (8 out of 25 chaperone interacting missense variants) and among variants of unknown significance (19 out of 32 chaperone interacting). Furthermore, BRCA1-BRCT missense variants that bind moderately to chaperones are equally prevalent in TCGA^67,68^ (somatically acquired or germline variants across diverse cancer types) as compared to the general population (Figure 4A). For 88 entries from TCGA, 24 variants moderately bound HSP70 (∼27%) as compared to 11 that strongly bound HSP70 (∼13%). Similarly, out of 84 germline BRCA1-BRCT variants catalogued in the gnomAD database^2^, 16 variants (∼19%) bind HSP70 moderately as compared to 8 variants (∼9.5%) that strongly interacted with HSP70. After, combining unknown significance BRCA1 variants from ClinVar, TCGA and gnomAD and filtering for redundant variants, ∼18% of variants moderately bound HSP70 as compared to ∼12% of variants that strongly bound HSP70 (Figure 4A). We observed no significant differences in the frequency of these mutation classes between different subpopulations (Figure 4B). In addition, missense variants observed in humans that confer hypomorphic phenotypes in cell-based variant screens more frequently bind moderately to HSP70 (∼32%, 9 out of 28 hypomorphic variants) as compared to loss-of-function variants which tend to bind strongly (∼58%, 29 out of 50) (Figure 4C)^50,52^. Thus, chaperones detect an affluence of hypomorphic BRCA1-BRCT variants in the general population and in tumors that may be relevant to human health.

**Figure 4.**
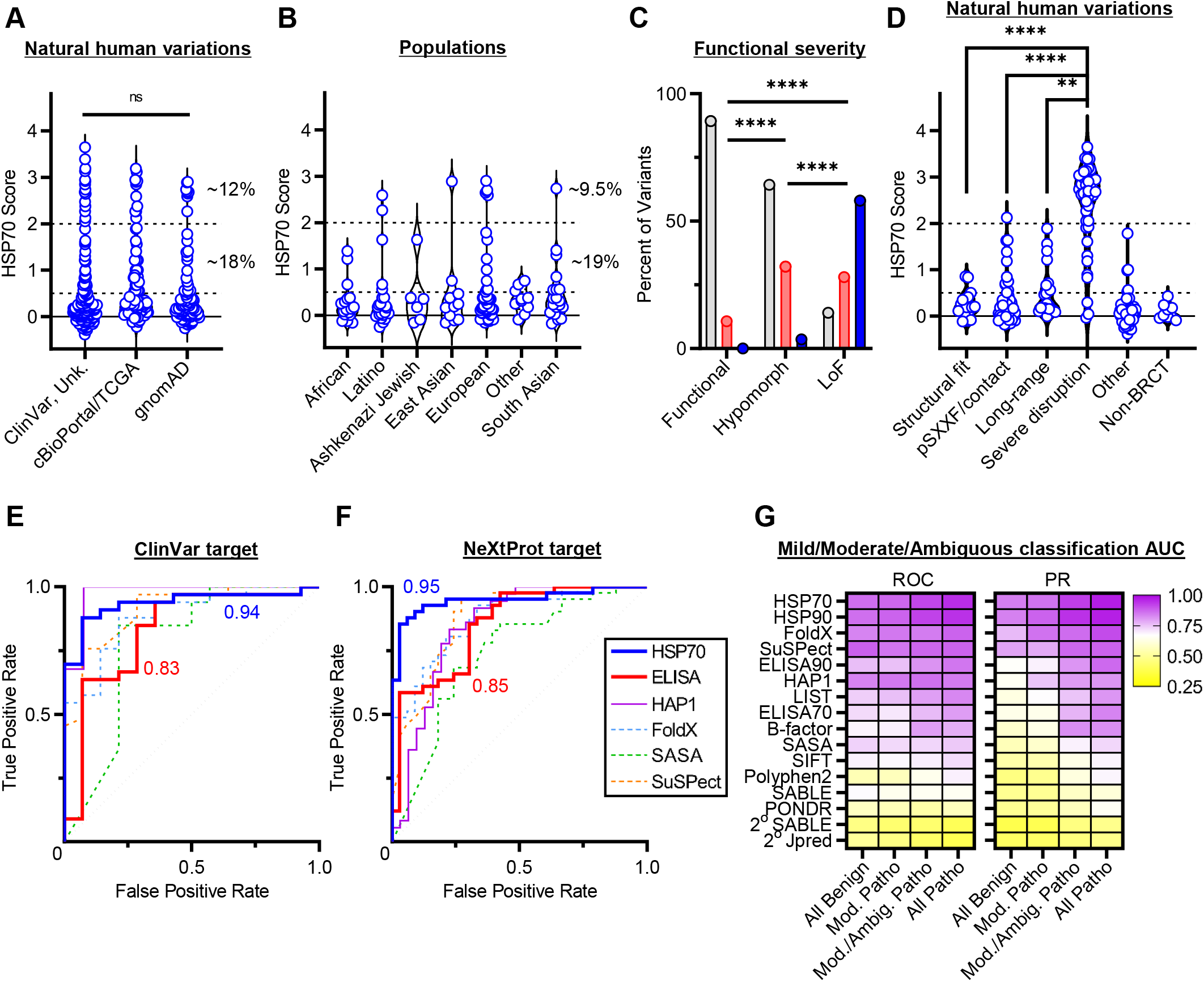
Chaperones identify diverse classes of pathogenic *BRCA1* variations in people. (A, B) HSP70 interaction scores to natural human variants observed in cancer patients (ClinVar^5^ and cBioPortal/TCGA^67,68^) or populations (gnomAD^2^). Dashed lines indicate cutoffs for moderate and strong HSP70 binding scores. Percentages derived by removing redundant variants such that each variant is counted only once. Unk., unknown. (C) Functional severity from cell-based variant screens^50,52^ of natural BRCA1-BRCT variants grouped by the degree of HSP70 binding. Gray bars show weak (HSP70<0.5), red bars show moderate (0.5<HSP70<2) and blue bars show strong HSP70 bound variants (HSP70>2). Data shown includes a non-redundant collection of variants from ClinVar, TCGA and gnomAD (n=176). Lof, loss of function. (D) HSP70 binding to natural BRCA1-BRCT human variants binned by the predicted effect on domain structure (see METHODS). (E, F) ROC curves using ClinVar or NeXtProt as the target dataset. NeXtProt curves assign mild/moderate/ambiguous variants as pathogenic. Area under the curve (AUC) for HSP70 (blue) and ELISA90 (red) parameters is shown. (G) AUC observed when mild/moderate/ambiguous variants are assigned differently in the NeXtProt target dataset. Statistical significance was determined using Kruskal-Wallis ANOVA test (A and D) or Chi-squared test (C). ****p≤0.0001, **p≤0.01. ns, no significance. Data are represented as mean from at least two independent experiments.

This observation motivated us to ask if hypomorphic *BRCA1* variations are subject to purifying selection in humans. We observed significant depletion of truncation *BRCA1* mutations from the general population as compared to missense variations, suggesting that loss-of-function *BRCA1* variants undergo purifying selection (Figure S4A, *p*≤0.0001). Interestingly, variants that bind moderately or strongly to chaperones exhibit allele frequencies similar to truncation mutants, suggesting that both variant groups are subject to purifying selection. However, differences in allele frequencies between variants that bind chaperones and those that do not bind chaperones were not statistically significant, likely due to the abundance of rare BRCA1 variants present in this population study (e.g., only 1 individual carrying a specific allele) and the possible influence of unaccounted risk factors. In contrast, many BRCA1 truncating mutations reach high allelic frequencies in tumors similar to missense mutations, suggesting reduced purifying selection of truncation mutants in the tumor (Figure S4B, *p*=0.02). It will be interesting to test how variants that bind moderately to chaperones contribute to structure-function relationships and cancer pathogenesis.

To better identify diverse classes of structure disrupting variants observed in humans, we asked if variants moderately bound by HSP70 share commonalities distinct from variants that severely disrupt BRCA1-BRCT structure. We parsed missense BRCA1 variants into structural effect classes to compare variants expected to partially disrupt domain structure with those expected to severely disrupt structure. We observed predominantly weak HSP70 binding to isosteric variants expected to be accommodated in the structure (e.g., mutating glutamates to glutamines) as compared to variants that severely disrupt structure (Figure 4D, structural fit *vs* severe disruption *p*≤0.0001). As noted above, we observed a number of variants within the hydrophobic pSXXF binding cleft that moderately bound HSP70 (10 out of 25), suggesting frequent disruption of the pSXXF binding cleft by complex mutations in people. Similarly, we observed weak to moderate HSP70 binding for variants that disrupt long-range interactions (side chain hydrogen bonds or inter-subdomain) as compared to severely disrupting variants (Figures 4D and S4C, long-range *vs* severe disruption *p*=0.002). Variants that we could not classify into discrete structural classes (‘other’) are largely surface variants that did not bind chaperones similar to variants outside of the BRCT domain (‘non-BRCT’). Many hypomorphic variants coincided with moderate increases in chaperone binding, predominantly mutating residues near the pSXXF and the inter-subdomain groove^50,52^ (Figure S4D). Interestingly, variants that interacted moderately with HSP70 were abundant in all structural effect classes we analyzed (excluding non-BRCT). Hence, classifying hypomorphic structure disrupting variants in people will be critical to identifying individuals carrying *BRCA1* alleles that increase cancer risk.

We tested the accuracy of chaperone interactions as indicators of pathogenicity as compared to other metrics standard in the field by utilizing receiver-operating characteristic (ROC) and precision-recall (PR) curves. Using pathogenicity annotated in ClinVar as the target dataset revealed near perfect accuracy when using HSP70 binding as the classifier (Figures 4E and S4E, area under the curve (AUC) ∼0.94). We also observed high accuracy when comparing ClinVar pathogenicity with variant function reported in HAP1^50^ (AUC ∼0.98) or variant stability predictions measured *in silico* (FoldX AUC ∼0.89), similar to the accuracy of chaperone binding. However, the performance of pathogenicity algorithms and cell function data decreased when we accounted for partial loss of function BRCA1 variants using NeXtProt as the target dataset. When mild/moderate/ambiguous functional severity variants were called pathogenic in the target data set, the accuracy decreased for most predictors (∼0.85 ROC AUC). In contrast, HSP70 binding reported near perfect accuracy in both ROC and PR curves across all pathogenicity assignments for mild/moderate/ambiguous variants (Figures 4F and S4F, AUC ∼0.95). Indeed, HSP70 and HSP90 binding outperformed variant expression levels as determined by ELISA (AUC ∼0.85), pathogenicity predictions by Polyphen2 or SuSPect, and biochemical metrics of the mutated residue such as solvent accessibility, regardless of how mild/moderate/ambiguous variants were assigned in the target dataset (Figures 4G and S4G). Notably, the accuracy of chaperone binding increased when variants with mild and moderate phenotypes were assigned into the pathogenic group, suggesting that chaperone binding is highly sensitive at detecting hypomorphic variants. Moreover, 50% (39 out of 78) of variants we tested from the NeXtProt server are either uncertain clinical significance or not annotated in ClinVar highlighting the potential of chaperone binding to assist in variant classification. Altogether, these results suggest that quantitative chaperone interactions can be used to identify pathogenic variants including hypomorphic alleles.

### Chaperone binding patterns stratify cancer risk and variant expressivity of human *BRCA1* carriers

*BRCA1* germline mutations have estimated breast cancer penetrance as high as 65%-85% and thus *BRCA1* mutation status is an actionable cancer predisposition marker in familial genotyping. However, significant variability in expressivity confounds predictions of cancer risk and age of onset. To determine if chaperone binding can elucidate the penetrance and expressivity of *BRCA1* mutations in carriers, we collected familial penetrance likelihood data of heritable *BRCA1* variants from the Leiden Open Variation Database (LOVD)^69^. Breast cancer risk was significantly increased for carriers of *BRCA1* mutations that increase HSP70 binding to the BRCA1-BRCT domain as compared to mutations that do not increase binding (Figure 5A, *p*≤0.03). ∼86% (18 out of 21) of BRCA1 variants that conferred increased breast cancer penetrance interacted with HSP70 (penetrance likelihood>1). Additionally, ∼67% (6 out of 9) of variants that interacted moderately with HSP70 conferred increased breast cancer penetrance. Furthermore, strongly HSP70-bound variants were highly penetrant for two independent penetrance parameters and the product likelihood ratio (Figures S5A-C), and we observed no significant association between the penetrance and strength of HSP70 binding. These results suggest that heritable hypomorphic BRCA1 variants that moderately bound HSP70 are important determinants of breast cancer risk. Interestingly, 4 highly penetrant variants that moderately bound HSP70 mutate residues near the pSXXF (D1692H, R1699W, M1775K and M1775R), suggesting that complex pSXXF cleft disruption drives increased penetrance in these carriers. In contrast, 2 highly penetrant variants moderately bound by HSP70 mutate buried hydrophobic residues that are not pSXXF coordinating (V1736A and M1783T) suggesting that partial disruption to domain folding drives increased penetrance. Hence, BRCA1-BRCT variants that compromise domain folding, even partially, can increase the likelihood of developing cancer.

**Figure 5.**
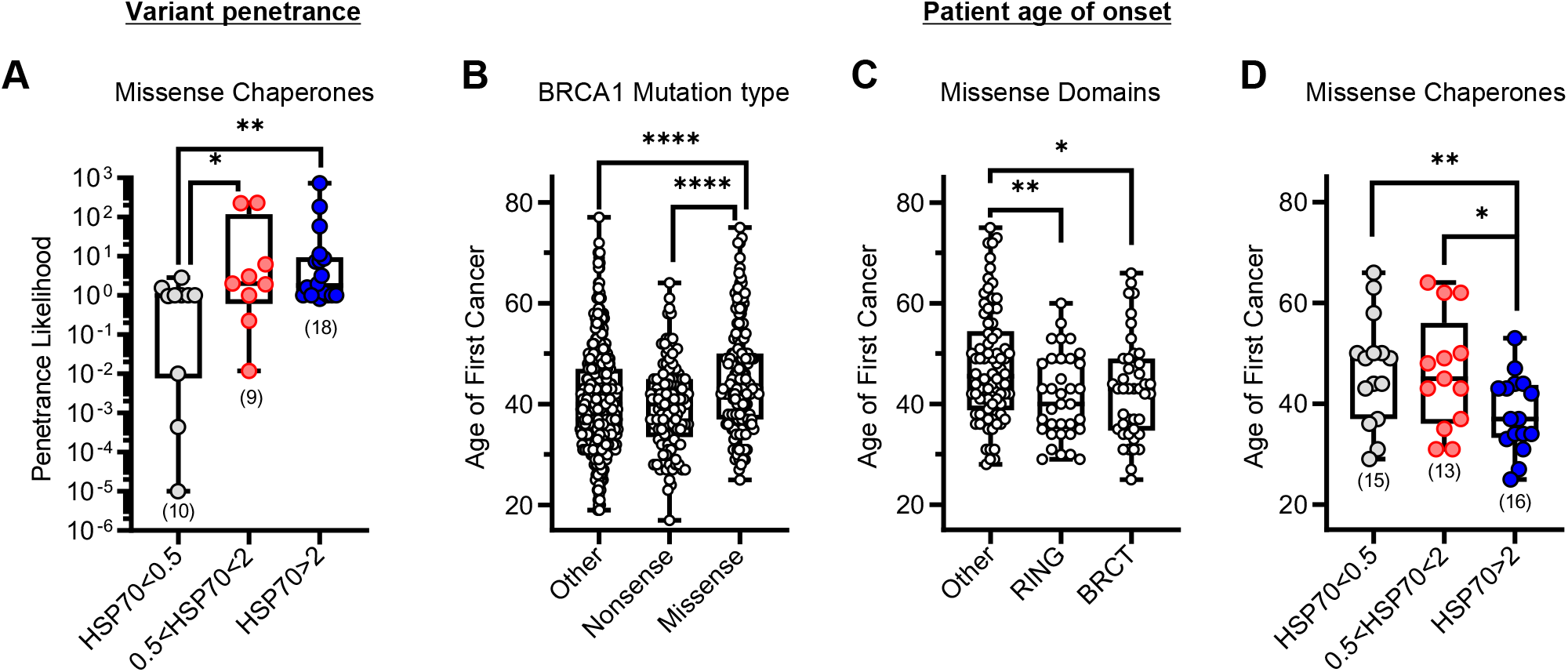
Chaperone binding patterns stratify cancer risk and variant expressivity of human *BRCA1* carriers. (A) *BRCA1* mutation penetrance likelihood binned by the magnitude of HSP70 binding. Data shown from the LOVD database^69^, accessed 4/24/2023. (B, C, D) Age of first cancer diagnosis for patients carrying *BRCA1* mutations binned by mutation type, missense mutations in domains and the degree of HSP70 binding. Statistical significance was determined using two-tailed (A) or one-tailed Mann-Whitney t-test (B, C and D). ****p≤0.0001, **p≤0.01, *p≤0.05. The number of variants in each bin shown in parentheses. Data are represented as mean from at least two independent experiments.

To validate our findings we evaluated an independent cohort of 1,106 *BRCA1* carriers^70,71^ who had clinical genetic testing at MD Anderson Cancer. 71% of individuals in this cohort developed breast cancer and the remainder were negative at the time of examination (Figure S5D). Comparing the genotypes of breast cancer patients and the ‘no cancer’ group did not reveal significant differences in sequence-based mutation types that could inform cancer risk predictions (Figure S5E). Thus, we asked whether increased chaperone binding to BRCA1 variants is associated with increased variant expressivity in this dataset. We observed that patients carrying a BRCA1 variant that binds to HSP70 had slightly reduced survival compared to patients carrying variants that bind weakly or moderately to HSP70 (Figure S5F, *p*=0.20). These results suggest that chaperone binding to variants carried in the germline can inform cancer prognosis by genotype. The low statistical significance of this endpoint analysis is likely due to the small number of deaths observable over 20 years. To improve our statistical power, we tested the relationship between increased chaperone binding and age of disease onset which was collected for each cancer patient at the time of diagnosis.

We hypothesized that individuals who develop cancer earlier in life carry a BRCA1 variant that is more functionally compromised, and thus, these individuals have a higher risk of developing cancer. Indeed, carriers of nonsense *BRCA1* mutations developed cancer significantly earlier than patients carrying missense mutations, consistent with this hypothesis (Figure 5B, *p*≤0.0001). Furthermore, missense mutations within the BRCT and RING domains resulted in earlier age of onset as compared to missense mutations in other regions of the BRCA1 protein (Figure 5C, *p*≤0.02). In accordance with our hypothesis, patients carrying BRCA1-BRCT variants that bind strongly to HSP70 developed cancer 12 years earlier (median age of onset) than patients carrying mutations that bind HSP70 weakly (Figure 5D, HSP70<0.5 *vs* HSP70>2: *p*=0.009). Strikingly, patients carrying BRCA1-BRCT variants that moderately bound HSP70 developed cancer with delayed onset as compared to carriers of strongly HSP70 bound variants (0.5<HSP70<2 *vs* HSP70>2: *p*=0.02, 8 year median delayed onset). FoldX predictions and HAP1 functional data also suggest that BRCA1-BRCT variants that partially disrupt structure and function manifest with delayed disease onset in the clinic (Figures S5G-H, between 8.5-11.5 year median delayed onset). These results show that hypomorphic, moderate HSP70 bound variants are an important, phenotypically distinct class of pathogenic *BRCA1* variations in humans. We conclude that the degree of chaperone binding can detect the penetrance and expressivity of cancer variations in people, enabling stratification of hypomorphic mutation carriers.

## DISCUSSION

Establishing the clinical significance of genetic variations relies on functional validation studies^72^. However, even for the well characterized *BRCA1* gene—a poster child for genetic testing—the fraction of useful variations that have been successfully integrated into clinical guidelines is dismal. This is neither due to an insufficient number of clinically relevant models^73,74^, nor the lack of high-throughput functional assays to evaluate variations *en masse*^50^. A major barrier to increasing the clinical relevance of genetic variation is linking the molecular mechanism of mutations to their clinical manifestations. For *BRCA1*, this barrier originates from the many roles BRCA1 plays within the cell, the complex effects of BRCA1 variations and the influence of numerous genetic and environmental modifiers, which contribute to incomplete penetrance and variable expressivity of *BRCA1* variations^6–8^. Despite remarkable advances in protein structure prediction and design, integrating data from functional assays and structure-predictions failed to lift this barrier for protein variants because the currently available solutions to the protein-folding problem are incomplete^24,25^. Here, we present a high-throughput approach to elucidate structure-function relationships based on measuring physical interactions between protein variants and ancient solutions to the protein folding problem, protein-folding chaperones. We discuss applications of this biochemical approach in precision medicine and variant structure prediction.

Protein-folding chaperones, including HSP70 and HSP90, recognize moieties nascent and misfolded polypeptides present on their surface transiently during folding^27,28^. The ability of chaperones to identify such transient (metastable) conformations is critical for making molecular decisions during protein quality control (i.e., forward folding, disposition, aggregation, degradation) and maintaining protein homeostasis within the cell. Chaperones evolved to serve these functions in the crowded intracellular milieu where the high frequency of collision between macromolecules shifts the protein-folding equilibrium towards aggregation. Chaperones, especially HSP90, can also modify the effects of disease-associated mutations^18,75–78^. We hypothesized that chaperones can be used as biosensors to infer the structural and phenotypic effect of mutations, thus bypassing the need for fully understanding structure-function relationships in the cell. Here, we demonstrate the validity of this hypothesis by using a quantitative high-throughput assay to measure the binding of HSP70 and HSP90 to >200 variants of the *BRCA1* tumor suppressor within living cells. Chaperone interactions correctly identified ∼77% of pathogenic variations in the BRCA1-BRCT domain from ClinVar, and 52 out of 62 times chaperones separated intolerant *vs* tolerant mutations at the same position (e.g., G1706E *vs* G1706A, T1685A *vs* T1685S); they also accurately predicted the severity of mutations, outperforming alternative established structure-function approaches.

Chaperones also enable classification of otherwise indistinguishable *BRCA1* mutations into discrete functional groups based on the pattern of chaperone binding to the encoded variant. We demonstrate that different HSP70 and HSP90 binding patterns to BRCA1 mutant proteins reflect differences in severity of the associated *BRCA1* mutation. Notably, we show that chaperone binding is proportional to both the structural and functional defect induced by mutation, establishing the link between mutational loss of function and chaperone binding: loss of protein folding. Even the absence of chaperone binding to variants is informative when accounting for available functional data, which enables identification of true separation-of-function contact mutants that selectively disrupt protein-partner interactions without affecting structure. Taken together, integration of chaperone interactions and functional data enabled classification of a diverse set of BRCA1-BRCT variants into the following functional groups: 1) loss of function variants that severely disrupt structure and function, 2) benign variants that fold similar to wild-type, 3) complex variants that interfere with both domain folding and protein-partner binding, 4) separation-of-function variants that specifically disrupt protein-partner interactions, and 5) hypomorphic variants that retain partial structure and function. Remarkably, humans carrying BRCA1 variants that increase chaperone binding exhibited increased cancer penetrance and expressivity, accentuating the utility of chaperone binding patterns as predictors of cancer risk in people.

The observations this work makes regarding *BRCA1* hypomorphs are particularly telling. Hypomorphic *BRCA1* alleles are known to confer intermediate phenotypes in mice and cancer cell lines^79–83^ as compared to *BRCA1* null^84^. While most *BRCA1* hypomorphs known were rationally designed (e.g., deletion of RING domain), natural *BRCA1* hypomorphs have been previously characterized and implicated in cancer predisposition (germline variants) and PARP inhibitor resistance (somatic mutations)^85–87^. One such natural *BRCA1* variant, R1699Q, confers intermediate cancer risk and Fanconi-Anemia-like symptoms^86,88–90^. Our data confirm R1699Q as a true separation-of-function BRCT variant that adopts wild-type-like structure, but is unable to recruit key DNA repair partner proteins harboring a pSXXF motif. Yet, such *BRCA1* separation-of-function mutations with intermediate phenotypes are rare in people. In contrast, we find that most BRCT mutations destabilize domain folding and drive protein degradation. This observation explains how BRCT mutations confer severe phenotypic effects; destabilization of the BRCT C-terminus drives full-length BRCA1 degradation, disrupting both BRCT-dependent and BRCT-independent roles of BRCA1 in genome maintenance. Moreover, our work reveals unexpected variation in severity among BRCA1-BRCT variants and identifies a new class of *BRCA1* hypomorphs. These *BRCA1* variants are prevalent in the general population and confer hypomorphic effects via a distinct mechanism: they fully support partner binding and genome maintenance but allow the encoded proteins to spend less time in functional conformations, thereby leading to intermediate phenotypic effects because of their metastability. We show that BRCA1-BRCT hypomorphs confer high-penetrance susceptibility to breast cancer with reduced disease expressivity, as determined by late age of onset. These findings identify a new class of *BRCA1* hypomorphic alleles with distinct clinical manifestations.

Our data may also have broad applicability for precision medicine and structure prediction approaches. BRCA1-BRCT variants can disrupt the ground states (thermodynamics), the transition states between conformations (kinetics), or both. Even for these important differences in structure disruption, increases in chaperone binding reflect disruption to structure in cells, regardless of the biophysical mechanism. To this point, our data provides a rich resource of chaperone binding data to BRCA1 variations that can be integrated with computational approaches and prediction algorithms based on machine learning to enhance predictions of biophysical mechanism driving structure disruption as well as variant assignment^24,25^. Notably, the high frequency of HSP70-engaged mutations in BRCA1-BRCT may indicate lower tolerance of the BRCT domain to mutations, as compared to the potentially higher mutational tolerance of the FANCA protein^18^. It would be interesting to test if HSP70-engaged BRCA1 variants hypersensitize tumors to genotoxic agents as compared to variants that are wild-type-like. Additionally, it would be interesting to test if variants that partially disrupt BRCA1-BRCT domain folding may be buffered by protein-folding chaperones, imparting an exquisite sensitivity to environmental stress for individuals carrying these variants. Interestingly, the disruptive effect of BRCA1 mutation can be rescued by HSP90, driving BRCA1-mediated cancer cell resistance^48^. Furthermore, chaperone interactions could be used to identify stable domains for biochemical and structural characterization and to facilitate protein design pipelines.

A limitation of this study is that the target variation set must map to globular domains to allow interrogation of changes in chaperone binding across different variants. >60% of residues in the human proteome are found in globular domains^12^, thus our approach has broad applicability for evaluating structure-function and genotype-phenotype relationships in humans. One can also envision instances where decreased chaperone binding to variants is informative, such as by binding of a co-factor or compound that induces protein stabilization, or a mutation that switches an intrinsically disordered protein to a well-structured protein. These ideas are easily tested with this approach. Altogether, in this study we highlight the utility of chaperone binding patterns as predictors of mutation severity and disease risk in humans.

## ACKNOWLEDGEMENTS

We thank Junjie Chen for providing plasmids encoding wild-type BRCA1 and for guidance with BRCA1 fragment expression. We thank Mikko Taipale for providing HEK293T cell lines stably transduced with lentiviruses constitutively expressing N-terminally NanoLuc-tagged HSP70 (*HSPA8*) and HSP90 (*HSP90AB1*) and Susan Lindquist for pcDNA3.1 destination vectors. We thank Natalia Condic for outstanding technical assistance, Nikitha Kota for help with ChaperISM predictions, Wenyi Wang and Shaolong Cao for help retrieving TCGA data, Jay Shendure and Greg Findlay for help accessing their HAP1 data and Richard Wood, John Tainer, Dan Dickinson, Swathi Arur, Pierre McCrea, Junjie Chen and Daniel Herschlag for feedback on data and the manuscript. Tumor allele frequencies are in whole based upon data generated by the TCGA Research Network: https://www.cancer.gov/tcga. GIK is a CPRIT Scholar in Cancer Research. Research reported in this publication was supported by the CPRIT under Award Number RR180005, and the National Cancer Institute of the National Institutes of Health under Award Number F32CA253780 to BG and K22CA222938 to GIK, from the Joe W. and Dorothy Dorsett Brown Foundation, the Robert J. Kleberg, Jr. and Helen C. Kleberg Foundation. The content is solely the responsibility of the authors and does not necessarily represent the official views of the National Institutes of Health.

## AUTHOR CONTRIBUTIONS

B.G. and G.I.K. conceived the project. B.G. designed and performed experiments with assistance from P.M. on LUMIER and coimmunoprecipitations. B.G. designed and constructed the BRCA1 variant library and performed computational analyses. A.G.B. and B.A. collected and curated the MD Anderson patient cohort with assistance from B.G. and G.I.K. B.G. and G.I.K. wrote the manuscript, with significant feedback and contribution from all other authors.

## MATERIALS AND METHODS

### Cell lines and culture

HEK293T stable cell lines expressing N-terminally tagged NanoLuc-chaperones (HSP70 (*HSPA8*) or HSP90 (*HSP90AB1*)) were maintained in 10-cm petri dishes (Corning, 08-772-22) grown in a humidity-controlled incubator at 37° C with 5% CO_2_ in DMEM (Corning, 10-013-CV) supplemented with 10%FBS (Sigma-Aldrich, F2442) and Pen Strep (Gibco, 15140-122). For passaging and experiments, we collected cells by treating with Accumax (Innovative Cell Technologies, AM-105), incubating at 37° C for 15-20 minutes and diluting with media. Cell lines were routinely tested for mycoplasma contamination (Lonza, LT07-318) and validated by STR fingerprinting by the MD Anderson Cytogenetics and Cell Authentication Core.

### Plasmids and Cloning

Plasmids containing human BRCA1 cDNA were a gift from Junjie Chen (Addgene, Plasmid #99394). We cloned the BRCT and RING variant library by designing primers (Sigma-Aldrich) to amplify the C-terminal (residues 1,314-1,863) or N-terminal (residues 1-324) region of BRCA1, flanked by attB handles compatible with Gateway® cloning. We purified PCR products by agarose gel extraction (Takara Bio, 740609) and used BP clonase (Invitrogen, 11789) to insert BRCT fragments into entry vector pDONR221. Then, we performed two-step PCR-mediated site-directed mutagenesis using Phusion HF (NEB, M0530) and primers containing the mutation of interest to generate mutant BRCA1 fragments flanked by attL sequences for LR clonase reaction (Invitrogen, 11791). Inserts were cloned into pcDNA3.1 expression vectors encoding tags (N-terminal: 3xFlag-V5, C-terminal: V5-3xFlag) and then propagated by transforming into DH5α bacteria and isolating single colonies on LB-plates supplemented with ampicillin. Plasmids were mini-prepped (Promega, A1222) and validated by restriction enzyme digestion and Sanger sequencing.

### LUMIER with BACON

LUMIER with bait control (BACON) was carried out as previously described using automated liquid handling robots (Biotek, EL406 and TECAN, Freedom EVO)^31,55^. Each experiment included eight replicates of the BRCA1-BRCT wild-type and four replicates of the RING wild-type plasmid DNA as well as controls to assess reproducibility. We arrayed 700 ng of plasmid DNA in round-bottom 96-well plates (Corning, 353227) and mixed with 200 µL PEIMax (Polysciences, 24765) with OptiMEM (Gibco, 319850) at ratio of 1:100 (10 µg/mL final). This DNA-transfection mix was then evenly distributed to four (two per HSP70 or HSP90 cell line) flat-bottom 96-well plates (Corning, 3598) and incubated at room temperature for 20 minutes. After, we reverse transfected by adding 100 µL of chaperone-luciferase fused stable cells to DNA-transfection mix in flat-bottom 96-well plates at a density of 30,000 – 50,000 cells/well in duplicate plates. Cells were grown for 2 days, washed three times with PBS using an automated Biotek plate washer and then incubated with lysis buffer (50 mM HEPES-KOH, pH 8.0, 150 mM NaCl, 10 mM MgCl_2_, 20 mM NaMo, 0.7% Triton X-100, 5% glycerol) supplemented with protease inhibitors, phosphatase inhibitors, RNAse and Benzonase for 5 minutes at room temperature. Lysates were transferred to 384-well plates pre-coated in-house with anti-FLAG antibodies (Sigma-Aldrich, F1804). Lysates were incubated at 4 °C for 3 hours with gentle rocking and then washed six times in cold HENG buffer (50 mM HEPES-KOH, pH 8.0, 150 mM NaCl, 2 mM EDTA, pH 8.0, 20 mM NaMo, 0.5% Triton X-100, 5% glycerol). 20 µL luciferase substrate was added to all wells at a 200x dilution for NanoLuc substrate (Promega, N11) and a 100x dilution for RenillaLuc substrate (Promega, E28) using reaction buffer (20 mM Tris HCl, pH 7.5, 1 mM EDTA, 150 mM KCl, 0.5% Tergitol NP9). Readings were taken using a 400-700 nm filter on an EnVision Plate Reader (Perkin-Elmer). For bait control, luciferase reagent was dumped into a sink and 30 µL of anti-FLAG coupled HRP (abcam, ab1238) diluted 1:10,000 in ELISA buffer (PBS, 5% Tween20, 1% goat serum (Sigma-Aldrich, G9023)) was added to all wells, and incubated for ninety minutes at room temperature with gentle rocking. Wells were washed six times with PBST (0.05% Tween20) and 30 µL ELISA Supersignal ELISA chemiluminescent substrate (Thermo Scientific, 370) was added and absorbance read.

### CoIP

Co-immunoprecipitations were carried out using agarose linked anti-FLAG beads (Sigma-Aldrich, F2426). 400,000 cells were seeded in 6-well plates on Day 0 (Corning, 3506). On Day 1, forward transfection was performed by mixing 3,200 ng of DNA packaged with 400 µL of OptiMEM combined with 8 µL of Lipofectamine2000, following incubation times as recommended by the manufacturer (Invitrogen, 11668). On day 3, cells were lysed with ice-cold LUMIER lysis buffer on ice for 20 minutes. Samples were then centrifuged at 14,000xg for 10 minutes at 4 °C, and the soluble fraction was mixed with pre-washed anti-FLAG beads and gently mixed for 3 hours at 4 °C. Beads were washed four times with ice-cold lysis buffer and proteins were eluted by adding 150 µg of 3xFlag peptide (Biomatik, 56305) competitor and gently rocking for 30 minutes at room temperature. Eluates were then collected and mixed 1:1 with a urea/SDS protein sample buffer followed by 65 °C incubation for 5 minutes.

### Western blotting

Denatured protein samples were loaded on pre-cast 4-12% SDS-PAGE gels (Invitrogen, WG1402) and resolved for 1.5 hours at 160 V in SDS-MOPS buffer (Invitrogen, NP0001). Proteins were transferred to PVDF membranes (BioRad, 1620177) by wet transfer using tris-glycine buffer supplemented with 10% fresh methanol. Membranes were blocked in 5% milk (Lab Scientific, M0841) prepared in PBST and incubated with corresponding antibodies described in key resources. Membranes were washed three times with PBST for 5 minutes each before incubating with anti-mouse secondary antibodies (Sigma-Aldrich, F1804) for at least 60 minutes, and then washed again before development using HRP substrate (Millipore, WBKLS0) and visualized using a Chemidoc MP (BioRad).

### Chaperone Interaction Scores

First, raw data was log_2_ transformed and each experiment was background subtracted by the average of non-transfected wells for each individual experiment. HSP70 and HSP90 stable cell line values were subtracted separately due to differences in the luminescence background signal of each cell line. We generated luminescence and ELISA z-scores by normalizing log_2_ transformed values to the log_2_ average and standard deviation of all non-transfected wells across all experiments. Then, ELISA z-scores were compared between experiments to filter data where the amount of bait protein purified varied from the median value obtained across all experiments (cutoff z-score<-2.5 or z-score>2.5, ∼1-2% of wells excluded). After these exclusions, log_2_ background subtracted values were averaged across all experiments to generate log_2_ signal to noise ratios (SNR). Variant chaperone interaction scores were generated by subtracting luminescence log_2_ SNR by ELISA log_2_ SNR to normalize for slight differences in abundance after pull-down, and then variant normalized values were subtracted by the average of all wild-type normalized values.

To determine chaperone interaction preference (HSP70 *vs* HSP90), we took the normalized chaperone interaction scores and converted them to interaction z-scores by normalizing HSP scores to the standard deviation of all HSP70 and HSP90 interaction scores. First, we identified variants that do not bind to either chaperones (‘none’, HSP interaction z-score<2.5). Then, variants that only bind HSP70 or HSP90 significantly were assigned HSP70>HSP90 or HSP90≥HSP70, respectively. For variants that exhibited increased binding to both HSP70 and HSP90, the ΔHSP interaction z-score (70 minus 90) was used to assign variants as HSP70>HSP90 (ΔHSP z-score>1) or HSP90≥HSP70 (ΔHSP z-score<1).

We binned BRCA1 missense variant data using the magnitude of chaperone binding to test if variants that bind HSP70 to a moderate degree reflect partial loss of function BRCA1 variants (we excluded frameshifting or truncating variants because the degree of chaperone binding cannot be directly compared with the wild-type BRCA1-BRCT due to change in length of the protein variant). In addition to the binning shown in the main text, we applied additional grouping strategies to test if the moderate chaperone binding variant group was robust to different binning approaches. Despite small differences in statistical significance, all our conclusions were robust to independent binning strategies, such as binning by HSP90 interaction magnitude (HSP90<0.5, weak; 0.5<HSP90<2, modest; HSP90>2, strong), HSP70 interaction quartiles (Q1/Q2, weak; Q3, modest; Q4, strong), luminescence signal z-scores (z-score<2.5, weak; 2.5<z-score<5, modest; z-score>5, strong), hybrid quartile and z-scores (Q1/Q2 (z-score<2.5), weak; Q3 (z-score>2.5), modest; Q4, strong) and various HSP70 interaction score cutoffs ((HSP70<1, weak; 1<HSP70<2, modest; HSP70>2, strong) (HSP70<1.3, weak; 1.3<HSP70<2.7, modest; HSP70>2.7, strong)).

### *In silico* Predictions of Variant Biochemical Properties

We predicted DnaK (bacterial HSP70) binding sites using ChaperISM^91^. The algorithm was run using the qualitative-mode prediction, and the number of binding sites predicted for variants were compared to the wild-type sequence. Variants were then grouped into three bins: unchanged (number of binding sites “stayed same”), more sites (“more binding sites” or “existing site became larger”), or less sites (“less binding sites” or “existing site became smaller”). We also predicted DnaK binding sites using LIMBO^92^ and observed no significant difference in chaperone binding when comparing variants in DnaK predicted sites relative to variants not within sites.

We used FoldX 5.0^20^ to build variant models and calculate stability of BRCT variants relative to the wild-type (PDB: 1t29). First, we repaired the wild-type structure using “RepairPDB”. Then, variant models were built with the “NumberofRuns” parameter set to 5 to derive averages and standard deviations. We extracted ΔΔG values relative to the corresponding wild-type model generated for each variant. Variants at G1738 are not shown in plots due to abnormally high ΔΔG values arising from modeling errors. The algorithm was run both in the presence and absence of the pSXXF for all variants to identify residues in proximity to the pSXXF in the crystal structure. Unless otherwise noted, all FoldX data shown are taken from predictions made without the pSXXF.

We collected mutation pathogenicity predictions from Polyphen2^22^ using the HumDiv classifier model, GRCh37/hg19 genome assembly and canonical transcript settings. SIFT^93^ predictions were made using the UniProt-SwissProt_2010_09 database, a median conservation score of 3.00 and sequences with more than 90% identity removed from predictions. LIST-S2^94^ was used to generate mutation pathogenicity predictions enhanced by taxonomy based sorting using the full-length BRCA1 sequence. SuSPect^95^ was used to collect variant pathogenicity predictions using the full-length BRCA1 sequence.

Secondary structure predictions were collected using the STRIDE web portal^96^ using the crystal structure repaired by FoldX (PDB: 1t29). We extracted B-factors for each amino acid using the swift.cmbi web portal again using the crystal structure repaired by FoldX. PONDR^97^ was used to calculate residue disorder using the full-length BRCA1 sequence and the VL-XT predictor. We collected solvent accessibility (SASA) values from the GetArea webserver^98^ using PDB files repaired by FoldX with radius of water probe set to 1.4 Å and without gradient calculations. We used SABLE^99^ to predict solvent accessibility and secondary structure using the full-length BRCA1 sequence with predictor type set to include Approximator (Exposure Pred). Secondary structure was also predicted by JPRED4^100^ using the 1314 to 1863 BRCA1 sequence.

### Selection and Characterization of Variants

*BRCA1* variants in ClinVar were extracted and curated to filter for single missense mutations. Variants were called pathogenic or benign including ‘likely’ classifications if the review status was at least ‘two’ gold stars. Variants we classified as ‘unknown’ in the text include entries that are annotated as uncertain significance, conflicting interpretations, not provided, or ‘one’ star review status. All ClinVar BRCT missense variants were tested using FoldX and variants were selected to represent benign and pathogenic variant groups. We cloned 11 out of 19 benign and 31 out of 56 pathogenic BRCA1-BRCT missense mutants from ClinVar. This includes a majority of the BRCA1-BRCT missense variants from ClinVar (56%) and thus our variant library is representative of the whole BRCT ClinVar missense mutant dataset. cBioPortal variants were extracted using the “curated set of non-redundant studies”. BRCA1 variants were extracted from 13 TCGA cancer types (KICH KIRP COAD BRCA LUAD LUSC PAAD PRAD LIHC HNSC BLCA KIRC READ). gnomAD variants were extracted from v2.1.1. Variants from cBioPortal, TCGA or gnomAD were filtered using FoldX to select for mutations predicted to either disrupt or have no effect on protein structure.

To test if chaperones detect changes in thermodynamic stability, a subset of ClinVar variants (n=44) were compared to previously published protease sensitivity data^47^. We generated a protease sensitive variant group using a <30% protease resistance cutoff (relative to wild-type). We compared these data to a control group of variants that are wild-type like in all assays (>70% cutoff). Variants that exhibited partial sensitivity to protease digestion or ambiguous functional effects were not included in these comparisons. 49 variants were rationally designed using BRCA1-BRCT crystal structures to disrupt or support protein folding (interactions we investigated were validated in five different BRCA1-BRCT crystal structures). ProteinTools^101^ was run on the BRCA1-BRCT PDB repaired by FoldX without the pSXXF to identify residues involved in hydrophobic networks, side chain hydrogen bond networks (including salt bridges) and long-range interactions (BRCT structure contact map). Residues in the crystal structure were defined as buried (SASA<10%), intermediate (10%<SASA<30%), or surface (SASA>30%). Surface folding determinants were defined as surface or intermediate variants (SASA>10%) that form side chain hydrogen bonds and bind HSP70 (HSP70 score>0.5), excluding proline variants introduced within α-helices which are expected to disrupt α-helix secondary structures.

We predicted whether variants would bind to chaperones by considering the biochemical properties of the wild-type residue and variant. Variants predicted to have structure disrupted and bind chaperones included 1) prolines in α-helices, 2) truncations within the BRCT domain, 3) buried variants (<10% SASA) introducing charged side chains, 4) buried variants that decrease hydrophobicity, 5) buried variants that introduce steric clashes, 6) hydrophobic network residues mutated to charged variants and 7) disruptions to side chain hydrogen bonding. Variants predicted to be structure unaffected were 1) BRCT deletion or mutations outside the BRCT domain, 2) hydrophobic swaps at buried positions, 3) isosteric or quasi-isosteric swaps at buried positions, 4) variants that maintain wild-type side chain hydrogen bonds and 5) all remaining surface variants (>10% SASA). Variants were considered to decrease hydrophobicity if they lost at least two carbon containing side chain groups. Buried variants were considered to create a steric clash when the side chain was increased by at least two functional groups. Variants were called isosteric if the mutant structure is the same as the wild-type (e.g., aspartate mutated to asparagine). Quasi-isosteric variants differed by one functional group.

We collected BRCA1 mutation data from high-throughput variant screens measured in HAP1^50^ or HeLa^52^ cell systems. Separation-of-function mutants were defined as variants that do not bind HSP70 (<0.5 interaction score) and exhibit intermediate or complete loss of function in either HAP1 cell fitness or HeLa HDR assays. Separation-of-function mutants were classified as ‘contact’ if they are pSXXF proximal or surface. Residues proximal to the pSXXF were identified using ΔSASA values comparing the presence and absence of the pSXXF (Δ>5%). Surface BRCT residues excluding those near the pSXXF were identified using SASA values calculated in the presence of the pSXXF (SASA>30%). We also extracted a large collection of BRCA1 functional data from the NeXtProt server^65^. Each functional assay classifies the effect of BRCA1 variants relative to wild-type as a “phenotype intensity” (No affect, mild, moderate, severe). We grouped the functional assays into eight categories: 1) aggregation/degradation, 2) cell localization, 3) cell viability, 4) DNA repair, 5) genome stability, 6) genotoxic stress, 7) protein-partner interaction and 8) transcriptional activation. For each category, we plotted chaperone binding binned by “phenotype intensity”. In our analysis, we included variants with multiple entries because they represent either a unique functional assay or publication for that variant. We generated a stacked bar graph by collecting all “phenotype intensity” results for variants in our library, splitting them into bins by the degree of chaperone binding and then calculating the percentage of alleles associated with each “phenotype intensity”. For this, we filtered out separation-of-function variants observed in HAP1 cell fitness and HeLa HDR assays and corresponding variants that mutate the wild-type residue of a separation-of-function variant (denoted as ‘inferred contact’) because these variants are frequently false negatives by chaperone binding (n=9 variants removed).

We classified human alleles as functional, hypomorphic, or severe using cell fitness and HDR cell-based variant data^50,52^ considering the following possibilities: 1) Functional in both assays, 2) Functional in one assay with no data available for the second assay, 3) Functional in one assay and intermediate in the second assay, 4) Functional in one assay and loss of function in the second assay, 5) Intermediate in both assays, 6) Intermediate in one assay and no data available for the second assay, 7) Intermediate in one assay and loss of function in the second assay, 8) loss of function in one assay and no data available in the second assay and 9) loss of function in both assays. Groups 1-2 were assigned “functional”, groups 3-6 were assigned “hypomorphic” and groups 7-9 were assigned “loss of function” (LoF). We tested for partially disrupted hypomorphic BRCA1 variants by binning variants into structural effect classes. Some variants can fit in more than one class, e.g. A1708P introduces a proline in an alpha helix but also disrupts inter-subdomain interactions. So, we designed a hierarchy to prevent variants from being grouped into multiple structural classes. Variants were grouped into structural classes using the following order: 1) protease sensitive, 2) pSXXF near / separation-of-function contact, 3) helix breaking prolines / charged variants in β-sheets / buried charged variants / buried loss of hydrophobicity, 4) inter-subdomain interactions, 5) hydrogen bonding disrupted, 6) buried hydrophobic swap / quasi-isosteric / isosteric swaps, 7) FoldX ΔΔG>5, 8) Non-BRCT and 9) Other. Residues participating in long-range interactions were defined as at least 50 amino acids apart in primary sequence and within 5Å in the BRCT contact map. Groups 1, 3 and 7 were combined into a ‘severe disruption’ bin, groups 4 and 5 were combined into a ‘long-range’ bin and group 6 was denoted ‘structural fit’.

### ROC/PR Analysis

We generated ROC and PR curves using the ROCR package in RStudio. Solvent accessibility and FoldX predictions were without the pSXXF. FoldX predictions for variants at G1738 were set to 20 kcal/mol to account for the abnormally high ΔΔG values arising from modeling errors (this improved FoldX parameter predictions). For diagnostics using the ELISA parameter, a minimum threshold was set to prevent outliers from skewing the results (this improved ELISA parameter predictions). Solvent accessibility (GetArea and SABLE), B-factor values, secondary structure predictions (both JPRED and SABLE) and disorder predictions (PONDR) are based on the wild-type BRCT residue and do not take into account the variant. Residues with lower solvent accessibility (more buried) or lower B-factor (more ordered) were considered to be pathogenic if mutated. For sequence-based secondary structure predictions (SABLE and JPRED4), α-helix and β-sheet predictions were summed to generate a parameter that reflects the probability of secondary structure at a given residue, and positions more likely to have structure were called pathogenic when mutated. The data were trimmed so that all variants have predictions available for all parameters tested (ClinVar target n=47, NeXtProt target n=74), excluding the HAP1 parameter which were run separately to avoid further trimming of the data. For HAP1 ROC/PR curves, a separate run was performed (ClinVar target n=44, NeXtProt target n=67). These separate runs did not affect the relative difference in accuracy between diagnostic parameters. For ROC/PR curve analysis using ClinVar as the ground truth, ‘one’ star review status entries were considered including ‘likely’ classifications to avoid data trimming. For ROC/PR curve analysis using aggregated data from the NeXtProt server as the ground truth, the mode functional consequence for each mutant was used to assign variants in the target data set as benign or pathogenic. For variants where the mode severity was mild, moderate, or multi-modal (‘ambiguous’), we ran multiple ROC/PR curves setting mild/moderate/ambiguous variants systematically to either benign or pathogenic as the ground truth.

### Patient Data Curation

Patients were identified utilizing a prospectively maintained database after institutional review board approval at The University of Texas MD Anderson Cancer Center. Informed consent was obtained from all human subjects. Genetic test results from 1996 to 2023 and the clinical characteristics from BRCA1 variants were included in this study. De-identified patient data including BRCA1 germline status was carefully curated to exclude ambiguous clinical data from the cohort. Genomic alterations reported by clinicians were cross referenced to the human BRCA1 sequence taken from NCBI to validate mutations. Genomic alteration in BRCA1 were then categorized as frameshift, gene deletion, in-frame deletion, intronic, large deletion, large duplication, large insertion, missense, nonsense, splice site, synonymous, or undeterminable (15 out of 1,121 individuals (1.3%) were undeterminable and filtered out). Kaplan-Meyer survivor curves were generated using an HSP70 score cut-off of 1.5 and artificial censorship at 20 years to observe patient survival probability on a reasonable time scale after cancer diagnosis. For patients that had multiple cancer incidences reported, the lowest age was taken as the age of first cancer diagnosis. When comparing age of first cancer between groups in the cohort, we utilized one-tailed Mann-Whitney (unpaired nonparametric) t-tests to specifically test for significantly decreased age of cancer onset. Most human subjects in this cohort are white females (∼49%) given that breast cancer is the predominant diagnosis among females with BRCA1 mutations and most relevant to their health. For our analysis using HSP70 binding and age of first cancer diagnosis, the racial diversity was much greater (∼2% Native American, ∼2% Other, ∼4% Asian, ∼23% African American, ∼30% Hispanic, ∼39% White) than the entirety of the cohort, suggesting that race does not substantially influence our chaperone binding age of onset results.

