## Supplemental Figures for "Protein-Folding Chaperones Predict Structure-Function Relationships and Cancer Risk in *BRCA1* Mutation Carriers"

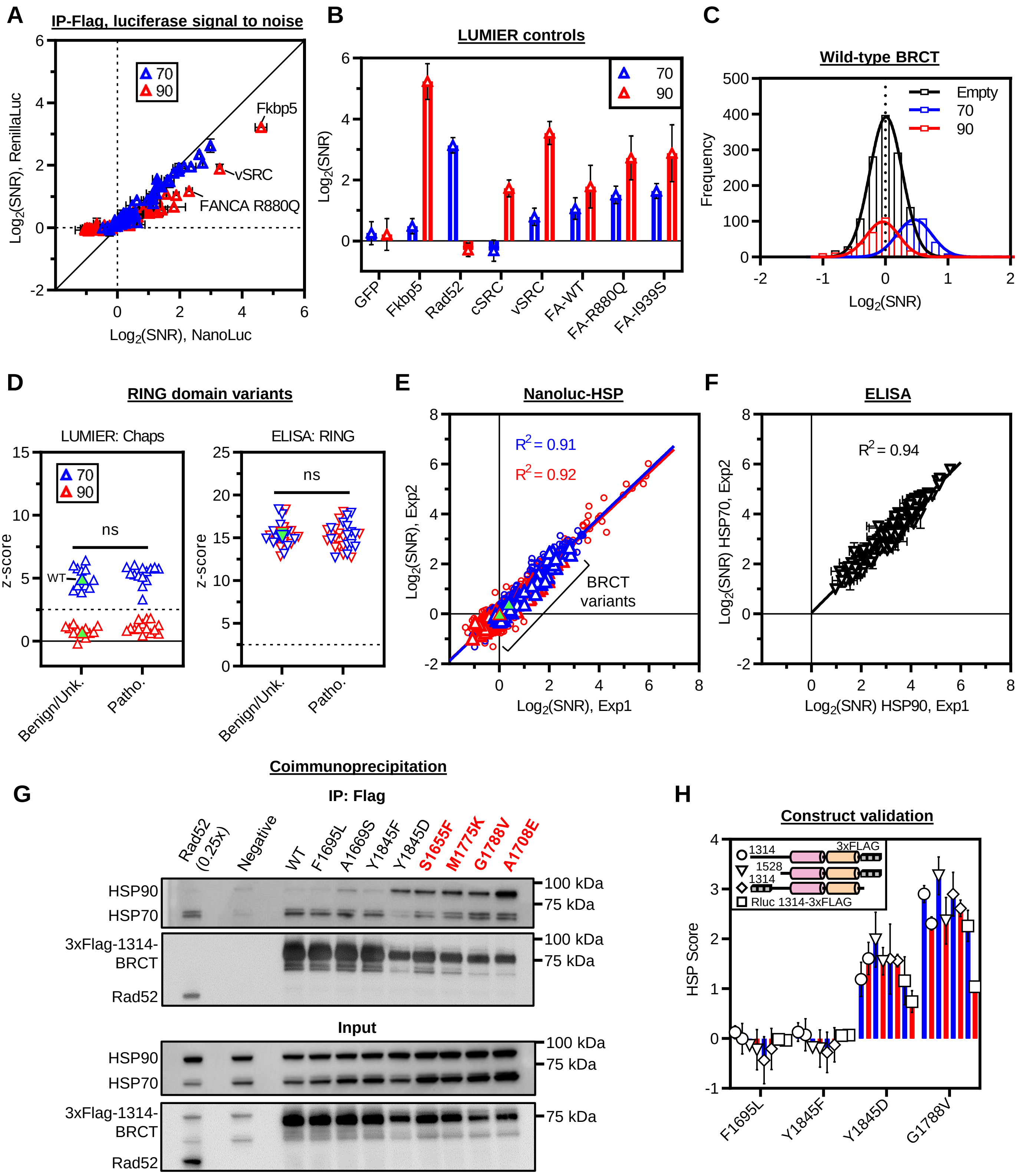


**Figure S1. BRCT variants induce chaperone binding to different degrees (related to Figure 1, legend on next page)**

**Figure S1. BRCT variants induce chaperone binding to different degrees (related to Figure 1)**

(A) Chaperone binding measured using LUMIER with NanoLuc (Nluc) or RenillaLuc (Rluc) cell lines. Signal to noise ratios (SNR) shown relative to negative control wells. Diagonal is identity line. All subsequent data is collected using Nluc cell lines unless otherwise noted.

(B) Chaperone binding to control proteins that exhibit known HSP70 and HSP90 binding patterns[18]. FA is FANCA proteins.

(C) Wild-type BRCA1-BRCT chaperone binding. Empty shows distribution of non-transfected control wells. HSP70 and HSP90 distributions show wells transfected with wild-type 1,314-BRCT-3xFLAG.

(D) Chaperone binding and ELISA measurements using BRCA1-RING variants annotated in ClinVar. The expressed RING fragment includes residues 1 to 324 tagged at the N-terminus with 3xFLAG. Z-scores are shown relative to non-transfected control wells. Wild-type (WT) RING values shown as green filled symbols. Dashed line indicates threshold for statistically significant signal (z-score>2.5). Benign and pathogenic groups are at least ‘one’ gold star review status, including ‘likely’ classifications. Unknown variants binned together with benign variants (unk.). Statistical significance was determined using two-tailed Mann-Whitney t-test. ns, no significance.

(E) Correlation of luminescence biological replicates quantifying chaperone binding. BRCA1-BRCT variants shown as triangles. Averaged wild-type BRCT values shown as green symbols for reference.

(F) Correlation of ELISA biological replicates quantifying bait-protein pull-down. Baits not detected significantly over background (z-score>2.5) excluded to quantify the correlation of detectable protein pull-down measured in different experiments.

(G) Anti-flag coupled agarose bead co-immunoprecipitation in HEK293T cells detected by Western Blotting against endogenous chaperones. Known pathogenic variants from ClinVar are shown in red. BRCT are N-terminally 3xFLAG tagged which gave greater signal than C-terminal tagged constructs. Rad52 is a positive control for HSP70 binding and loaded at 0.25x to account for strong HSP70 binding to Rad52 in the IP fraction.

(H) Variant chaperone interaction scores relative to wild-type in different truncations, tags, and Rluc reporter cell lines. Data are represented as mean±standard deviation from at least two independent experiments.

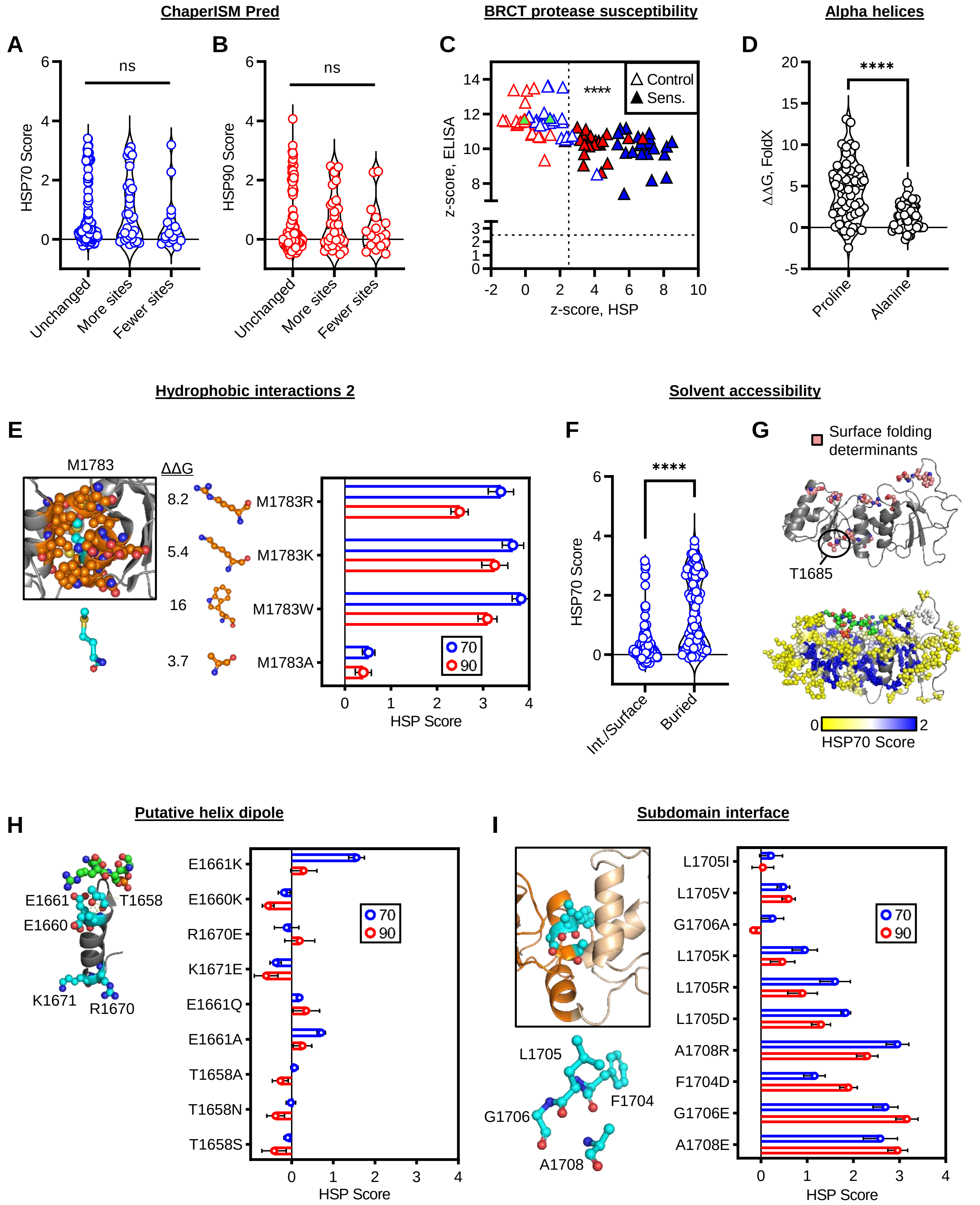


**Figure S2. Chaperones bind BRCA1-BRCT variants that disrupt structure (related to Figure 2, legend on next page)**

**Figure S2. Chaperones bind BRCA1-BRCT variants that disrupt structure (related to Figure 2)**
(A, B) Chaperone binding to BRCA1-BRCT variants grouped by number of DnaK sites predicted by ChaperISM[91]. ClinVar variant library.
(C) Chaperone binding and ELISA z-scores for BRCA1-BRCT variants previously characterized by protease digestion[47]. Filled symbols show variants sensitive (sens.) to protease digestion. Wild-type values are shown as filled green symbols.
(D) FoldX predictions from proline or alanine mutational scanning of residues in α-helices.
(E) M1783 variants to disrupt or support hydrophobic interactions. FoldX ΔΔG predictions for each variant are shown.
(F) HSP70 binding binned by solvent-accessibility. 10% accessibility cut-off computed without the pSXXF. Int., intermediate. ClinVar variant and rationally designed libraries.
(G) Chaperone binding to variants overlaid onto the crystal structure. Top, BRCT surface folding determinants ascertained by identifying surface variants that bind HSP70 and disrupt side chain hydrogen bonds. Bottom, HSP70 binding scores overlaid on the structure. For positions with multiple variations, the highest HSP70 binding score is shown.
(H, I) Rationally designed variants targeting interactions at α-helix 1 (H), and disruption of the subdomain interface between BRCT1 and BRCT2 (I). Statistical significance was determined using Kruskal-Wallis ANOVA test (A, B) or two-tailed Mann-Whitney t-test (C, D and F). ****$p \leq 0.0001$. ns, no significance. Data are represented as mean±standard deviation from at least two independent experiments.

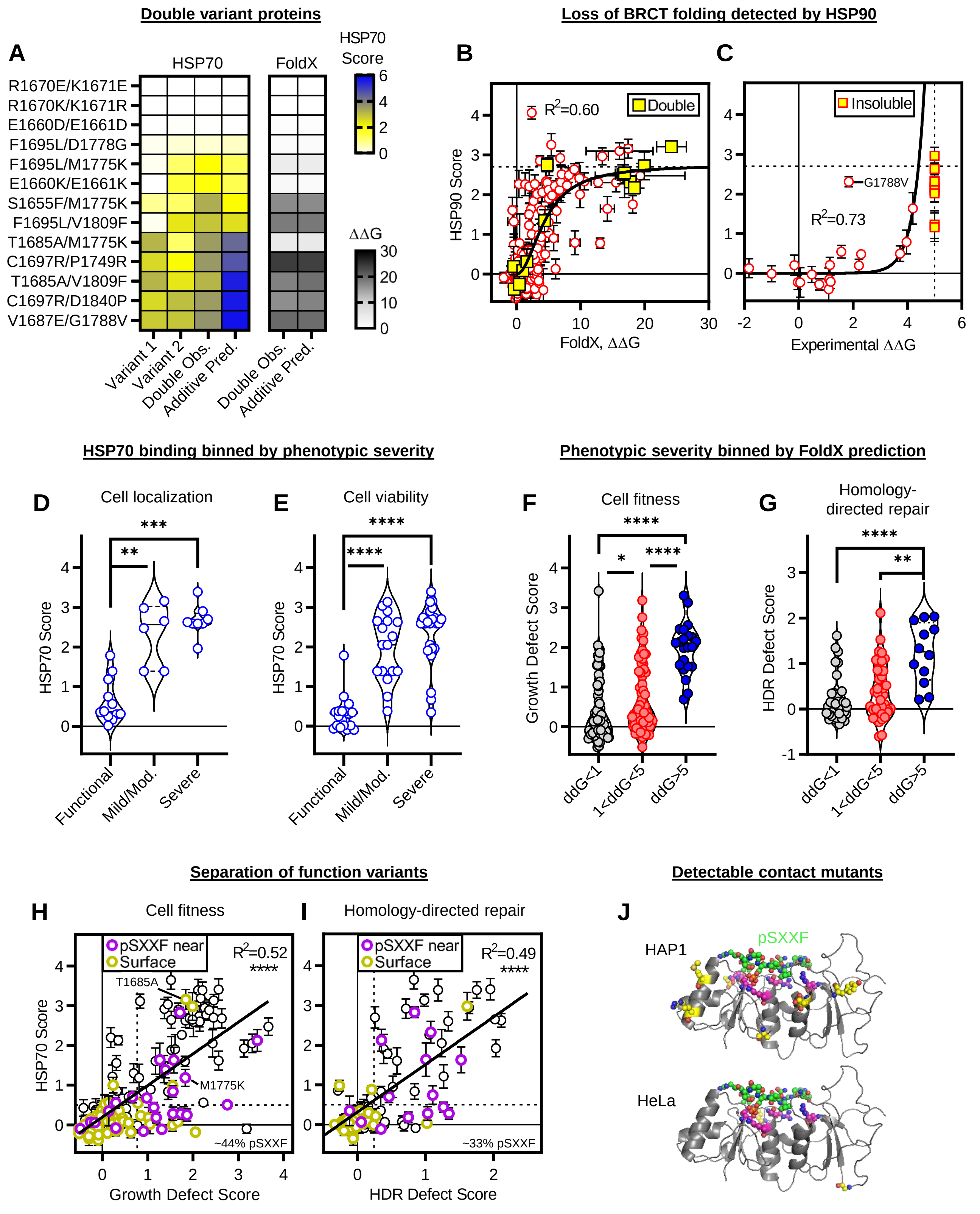


**Figure S3. Chaperone interactions delineate specific BRCA1 variant structure-function outcomes in cells (related to Figure 3, legend on next page)**

**Figure S3. Chaperone interactions delineate specific BRCA1 variant structure-function outcomes in cells (related to Figure 3)**

(A) HSP70 binding and FoldX predictions to single and double variants. Double observed (obs.) column shows chaperone binding measured for double variants. Additive predictions (pred.) column shows expected binding calculated by multiplying the fold increase in binding observed for both single variants.

(B, C) HSP90 binding correlated with protein folding stability. Horizontal dashed line is the average of four double variants that combine strongly chaperone bound single variants. Vertical dashed line in (C) reflects an upper limit due to variant insolubility. FoldX data in (B) fit to the hill equation with hill coefficient set to 2 to account for HSP90 dimerization. Experimental data in (C) fit to a single exponential growth (excluding insoluble variants). G1788V not fit because stability measurements were previously reported as contradictory[64].

(D, E) HSP70 binding to BRCA1-BRCT variants binned by the functional effect measured in cell localization and cell viability assays[65]. A majority of cell localization and viability assays were carried out using the full-length BRCA1. The same variant may appear more than once if the functional assay or reporting publication were different.

(F, G) BRCA1 variant function in HAP1[50] (cell fitness) or HeLa[52] (homology-directed repair) cells binned by FoldX prediction.

(H, I) Allele defect in HAP1 (cell fitness) or HeLa (homology-directed repair) cells correlated with HSP70 binding to the encoded variant protein. pSXXF near variants (purple) determined using ΔSASA when pSXXF is removed as compared to the presence of the pSXXF (>5%), and surface variants (mustard) was determined using SASA with pSXXF present (>30%). Vertical dashed line shows the cutoff for functional effect in cells (intermediate and loss of function) and horizontal dashed line shows cutoff for increased chaperone binding (HSP70>0.5). Separation-of-function variants that are functionally defective and do not bind chaperones are within the bottom right quadrant enclosed by the dashed lines. The percentage of separation-of-function variants near the pSXXF is shown in the bottom right corner of each plot. Data reproduced from Figures 3F-G to illustrate separation-of-function variants and the strong correlation between phenotypic effect and chaperone binding.

(J) Crystal structures highlighting separation-of-function contact variants determined by comparing cell functional data and chaperone binding (pSXXF near, magenta; surface, yellow). PDB: 1t29. Statistical significance was determined using Kruskal-Wallis ANOVA test (D, E, F and G). ****$p \leq 0.0001$, ***$p \leq 0.001$, **$p \leq 0.01$, *$p \leq 0.05$. Data are represented as mean ± standard deviation from at least two independent experiments.

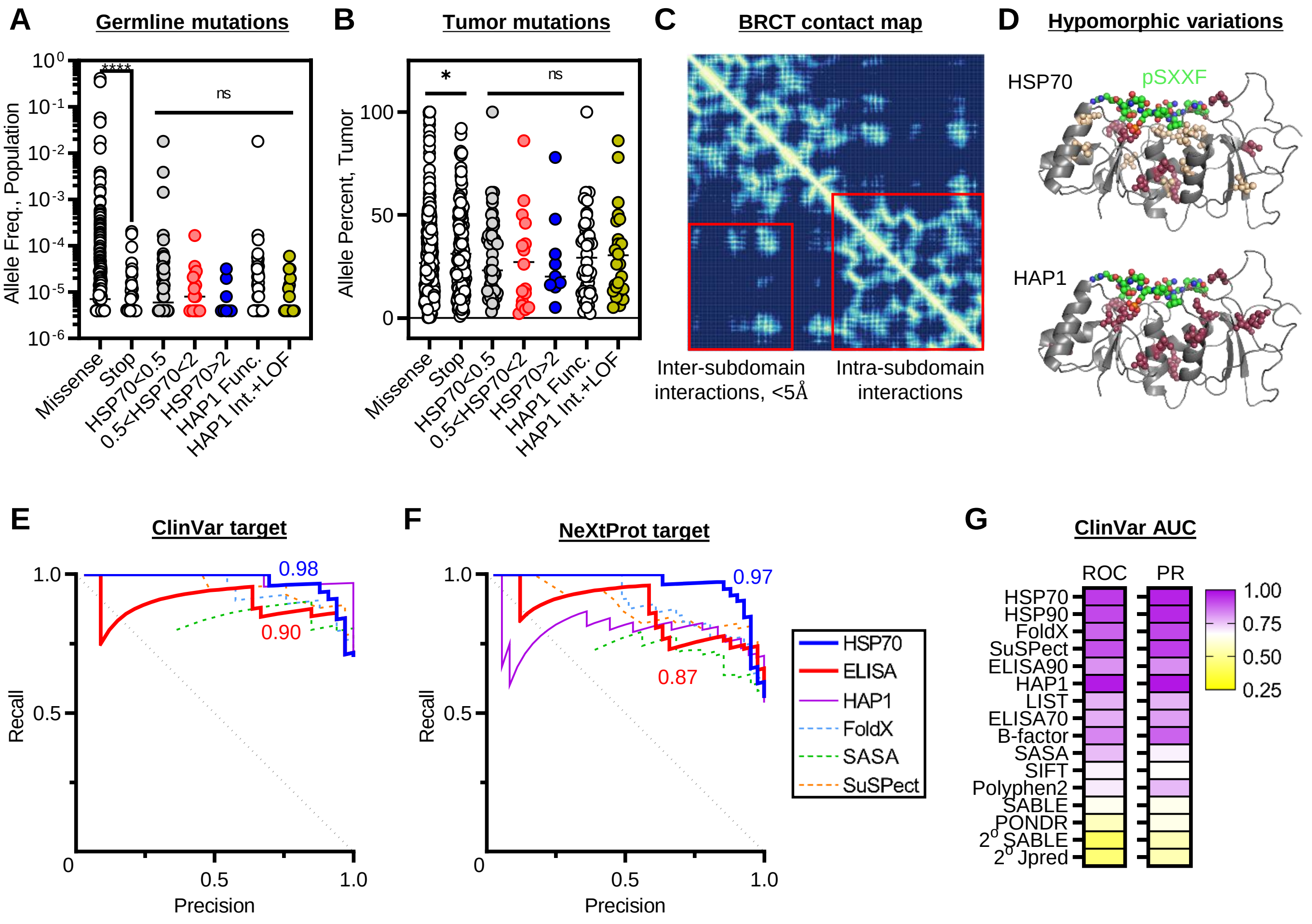


**Figure S4. Diverse features of human hypomorphic BRCA1 variants accurately identified by chaperones (related to Figure 4)**

(A, B) *BRCA1* allele frequencies from germline mutations observed in gnomAD (A) or tumor mutations observed in cBioPortal (B). Stop bin includes frameshifting and nonsense variants. Statistical significance was determined using two-tailed Mann-Whitney t-test. ****$p \leq 0.0001$, *$p \leq 0.01$. ns, no significance.

(C) BRCT crystal structure contact map. Lighter regions indicate amino acids close in three-dimensional space. PDB: 1t29 without the pSXXF.

(D) Moderate HSP70 bound variants overlaid on the BRCT crystal structure. Moderate HSP70 bound variants that overlap with hypomorphic *BRCA1* mutations observed in HAP1 variant screens[50] colored raspberry. Putative hypomorphic variants detected by HSP70 are colored wheat.

(E, F) Precision-recall curves using ClinVar or NeXtProt as the target dataset. NeXtProt curves assign mild/moderate/ambiguous variants as pathogenic. AUC for HSP70 (blue) and ELISA90 (red) parameters shown is shown.

(G) AUC for ROC and PR curves using ClinVar as the target data set.

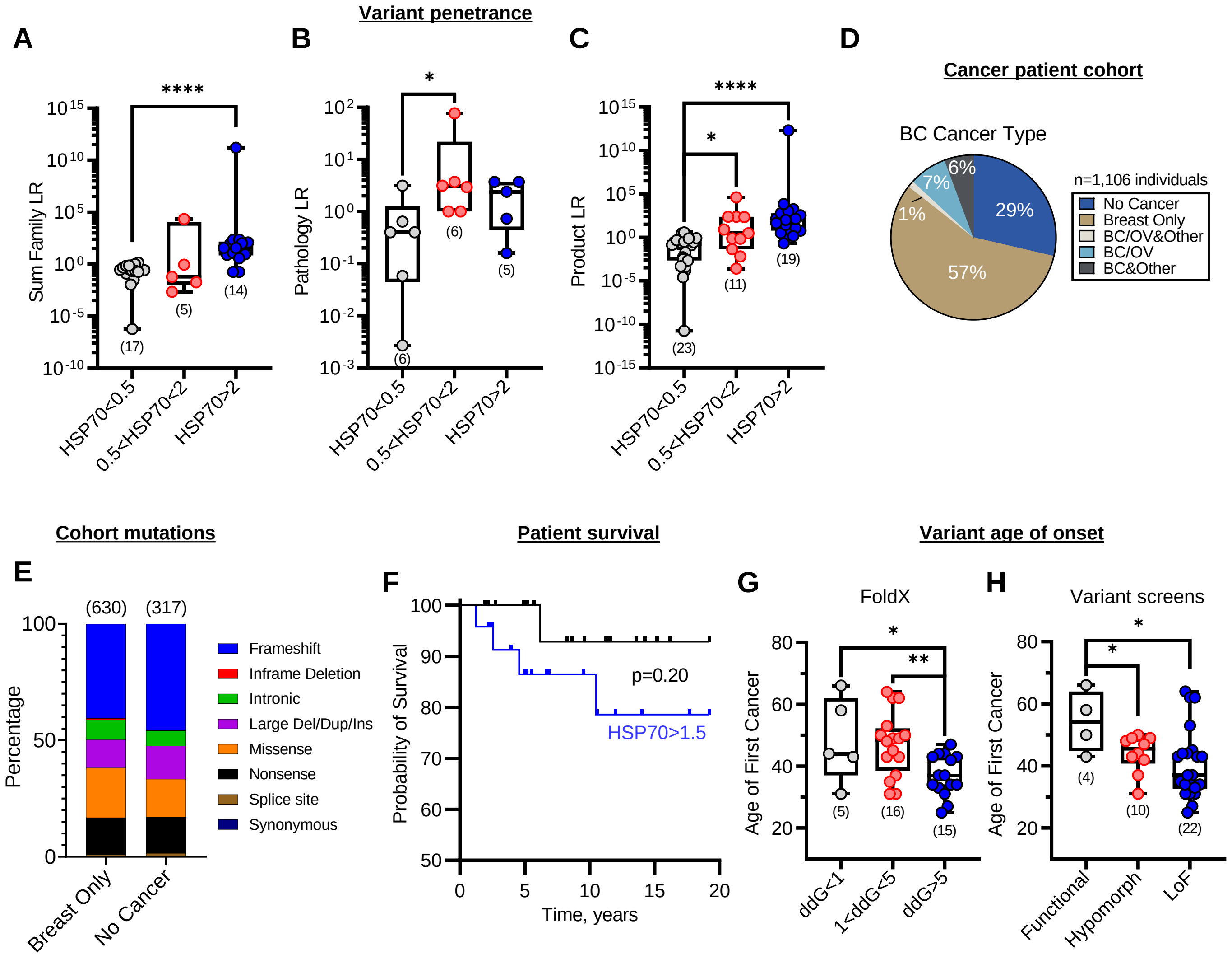


**Figure S5. Chaperone binding elucidates mutation penetrance and expressivity in human *BRCA1* carriers (related to Figure 5)**

(A, B, C) *BRCA1* mutation penetrance likelihood ratios (LR) binned by the magnitude of HSP70 binding. Data shown from the LOVD database[69], accessed 4/24/2023.

(D, E) Prospective MD Anderson human cancer patient cohort. Pie chart in (D) shows cancer type and bar chart in (E) shows *BRCA1* genotypes for breast cancer only and no cancer control group. Groups not visible in (E) are <0.4%.

(F) Kaplan-Meier survival curves for patients carrying germline BRCA1 variants that bind chaperones. Patients were artificially censored at 20 years to test survivorship on a reasonable time scale following cancer diagnosis. Black curve (HSP70<1.5), n=20. Blue curve (HSP70>1.5), n=24.

(G, H) Age of first cancer diagnosis for patients carrying *BRCA1* mutations binned by FoldX prediction or functional effect measured in HAP1 cells[50]. Statistical significance was determined using two-tailed Mann-Whitney t-test (A, B and C), Gehan-Breslow-Wilcoxon test (F), or one-tailed Mann-Whitney t-test (G and H). ****$p \leq 0.0001$, **$p \leq 0.01$, *$p \leq 0.05$. The number of variants in each bin are shown in parentheses. Data are represented as mean from at least two independent experiments.
